# Cerebral cavernous malformation 1 determines YAP/TAZ signaling dependent metastatic hallmarks of prostate cancer cells

**DOI:** 10.1101/2020.04.23.057778

**Authors:** Sangryoung Park, Ho-Yong Lee, Hansol Park, Young Seok Ju, Jayoung Kim, Eung-Gook Kim, Jaehong Kim

**Affiliations:** Department of Biochemistry, College of Medicine, Gachon University, Incheon 21999, Republic of Korea; Department of Health Sciences and Technology, Gachon Advanced Institute for Health Science and Technology, Gachon University, Incheon 21999, Republic of Korea; Biomedical Science and Engineering Interdisciplinary Program, Korea Advanced Institute of Science and Technology, Daejeon 34141, Republic of Korea; Graduate School of Medical Science and Engineering,, Korea Advanced Institute of Science and Technology, Daejeon 34141, Republic of Korea; Division of Cancer Biology and Therapeutics, Departments of Surgery & Biomedical Sciences, Samuel Oschin Comprehensive Cancer Institute, Cedars-Sinai Medical Center, Los Angeles, CA, USA; Department of Biochemistry, Chungbuk National University College of Medicine, Cheongju 28644, Republic of Korea

**Keywords:** Cerebral cavernous malformation, prostate cancer, metastasis, YAP/TAZ signaling

## Abstract

Enhanced Yes-associated protein (YAP)/transcriptional co-activator with PDZ-binding motif (TAZ) signaling is correlated with the extraprostatic extension of prostate cancer. However, the mechanism by which YAP/TAZ signaling becomes hyperactive and drives prostate cancer progression is currently unclear. In this study, we demonstrated that CCM1 induces the metastasis of multiple types of prostate cancer cells by regulating YAP/TAZ signaling. Mechanistically, CCM1, a gene mutated in cerebral cavernous malformation, suppresses DDX5, which regulates the PLK1-mediated suppression of YAP/TAZ signaling, indicating that CCM1 and DDX5 are novel upstream regulators of YAP/TAZ signaling. We also revealed that higher expression of CCM1, which is uniquely found in advanced prostate cancer, is inversely correlated with metastasis-free and overall survival in patients with prostate cancer. Our findings highlight the importance of CCM1-DDX5-PLK1-YAP/TAZ signaling in the metastasis of prostate cancer cells.

**Statement of Significance:** Our analysis of CCM1 expression and function represents a candidate predictive biomarker for prostate cancer metastasis and provides an evidence that abnormality of CCM1 can be pathogenic in prostate cancer. Importantly, CCM1 regulation of metastasis progression appears to a common molecular event in metastatic prostate cancer cells arising in disparate genetic backgrounds.

## Introduction

Prostate cancer (PCa) is a clinically heterogeneous disease with marked variability in patient outcomes (1-3). Importantly, PCa is also the second most common cause of male cancer death worldwide. Indeed, PCa is a disease entity for which no effective therapies exist once it progresses to metastatic castration-resistant PCa (mCRPC), and clinically advanced PCa is responsible for more than 250,000 deaths worldwide annually (4,5).

Advancements in therapeutic strategies for PCa have led to a surge in the incidence of metastasis, indicating the importance of preventing these detrimental metastatic events (6,7). Diverse sets of cancer hallmarks associated with aberrant functioning of the androgen receptor (AR), which is induced by androgen deprivation therapy (ADT), are the key driving forces behind the uncontrollable growth and metastasis of PCa and its transition into mCRPC (8-10). However, the clinically heterogeneous, multifocal nature of PCa along with the late appearance of castration resistance (CR) and the resultant uncontrollable bone metastasis makes the disease frustratingly complex to investigate and treat (10,11). Therefore, identifying the central molecular changes caused by current ADT-based therapeutic interventions, ultimately leading to the acquisition of uncontrollable metastasis, is critical for gaining important insights into key unmet needs, namely the improvement of diagnostic biomarkers and therapeutic interventions for mCRPC.

Yes-associated protein (YAP) and transcriptional co-activator with PDZ-binding motif (TAZ) signaling pathways have emerged as important drivers of the development, growth, and metastasis of human malignancies including PCa, and accumulating evidence has illustrated that YAP/TAZ signaling is correlated with the metastasis of PCa (12,13). YAP is a transcriptional co-activator that interacts with the transcription factor TEAD, and the interaction between YAP and TEAD is crucial for the expression of YAP target genes. Promoter regions of YAP/TAZ target genes have TEAD binding sites, and the interaction of YAP or TAZ with a member of the TEAD family of transcription factors increases TEAD transcriptional activity.

Increased expression of YAP or TAZ or their nuclear localization is observed significantly more frequently in mCRPC than in primary PCa, and disruption of YAP/TAZ or AR signaling suppresses the castration-resistant growth, motility, and invasion of PCa cells (12-15). Elevated YAP/TAZ signaling is also a positive regulator of AR signaling, and it responsible for CR and metastasis, indicating the significance of YAP/TAZ signaling in PCa progression (12,15). However, the mechanism by which YAP/TAZ signaling becomes hyperactive and drives PCa progression is currently unclear. In this study, we sought to clarify the novel regulatory mechanism of YAP/TAZ signaling in PCa progression using multiple types of PCa cells.

## Results

### 1. Higher expression of CCM1 at the mCRPC stage and its association with poor prognosis of patients with PCa

First, we analyzed changes in the expression of cerebral cavernous malformation 1 (CCM1) gene during PCa progression using data from multiple human PCa cohort studies. From the GDS2545 and GDS2547 datasets (16), we observed that CCM1 expression was remarkably increased in mCRPC samples (Figure 1A–B). Next, we applied a larger dataset from the PCa transcriptome atlas (PCTA), which we recently built using integrated bioinformatics analysis methods (17). From the PCTA, we also observed that CCM1 levels were dramatically increased only in samples from patients with mCRPC compared with those in primary prostate tissue and other stages of PCa (Figures 1C–D and S1A–C). The stem plots with different colors in Figure 1C illustrate that average CCM1 level was uniquely increased in mCRPC. The waterfall plot in Figure 1D displays normalized CCM1 gene expression for individual samples. We further examined the expression of CCM2, another member of the CCM family. Contrary to the findings for CCM1, we observed no meaningful increases of CCM2 or CCM3 expression during PCa progression (Figures 1E and S1D–F). The specific increase of CCM1 expression in the advanced stage of PCa suggests its potential as an index of poor prognosis for patients with PCa. We therefore analyzed the associations of CCM1 expression with overall survival (OS) in the Swedish Watchful Waiting cohort (18) and with metastasis-free survival in the Johns Hopkins cohort (19) (Figure 1F). We observed inverse correlations of CCM1 expression with OS and metastasis-free survival. Our analysis illustrated that increased CCM1 expression was significantly associated with poor prognosis for patients with PCa.

**Figure 1.**
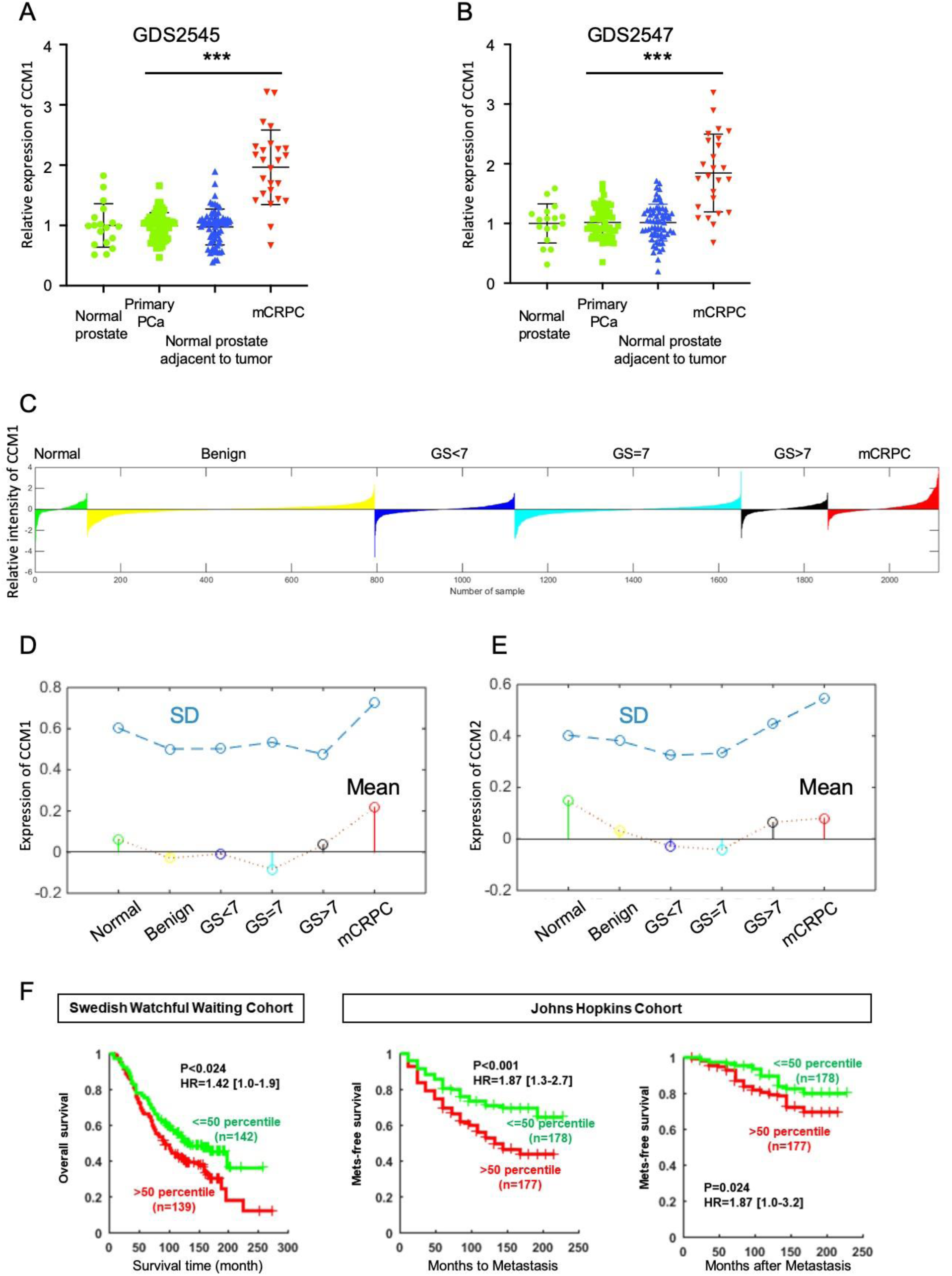
CCM1 levels are increased exclusively in metastatic castration-resistant prostate cancer (mCRPC) samples. (A, B) Relative CCM1 mRNA levels in normal prostate tissue (n = 17), normal prostate tissue adjacent to the tumor (n = 58), and primary (n = 64) and mCRPC (n = 25) tumors. Data were retrieved from accession numbers GDS2545 (A) and GDS2547 (B) of the Gene Expression Omnibus database. Data are presented as the mean ± SD. ***, P < 0.0001 (*t*-test). (C) This waterfall plot displays normalized CCM1 gene expression levels in individual samples from the prostate cancer transcriptome atlas (PCTA) cohort (n = 2115), which includes normal (n = 121), benign (n = 673), primary prostate cancer (PCa) with Gleason Sum (GS) < 7 (n = 328), GS = 7 (n = 530), or GS > 7 (n = 203), and CRPC/Met samples (n = 260). The x-axis presents samples, and the y-axis presents normalized gene expression (log2 scale). The Wilcoxon rank sum test results between subsets (mCRPC vs. primary PCa): Fold change = 0.256, P < 0.001. (D) The stem plots with different colors show the mean expression levels of CCM1 in each disease state from the PCTA cohort. The x-axis presents the disease state, and the y-axis presents normalized gene expression. The dot plot above the stem plot represents the variance (SD) of gene expression values of the samples from the same disease state. (E) The stem plots with different colors show the mean expression levels of CCM2 in each disease state. (F) To assess the association of CCM1 gene expression with overall or metastasis-free survival, clinical information was extracted from the Swedish Watchful Waiting Cohort (left) and Johns Hopkins Cohort (right two panels). Patients in each cohort were subdivided into “low” (<50th percentile) or “high” (≥50th percentile) CCM1 expression groups. Kaplan–Meier curves for overall and metastasis-free survival were drawn for each category, and Cox proportional hazard regression analysis was performed for statistical comparisons of survival rates between the low and high expression groups.

### 2. CCM1 induces metastatic hallmarks of PCa cells

Different PCa cell models possess distinct tumorigenic subpopulations that both commonly and uniquely express important signaling pathways (20). Considering the potential regulatory function of CCM1 in metastasis, we examined its involvement in metastasis-related features in multiple types of metastatic PCa cells. We generated a series of shCCM1 cell lines (PC3, DU145, LNCaP, C4-2, C4-2B, and CWR22r) in which CCM1 expression was stably suppressed using RNAi. We investigated changes in migration and invasion following CCM1 suppression using wound-healing assay and Transwell migration and invasion assays. Both PC3 and DU145 shCCM1 cells exhibited delayed wound closure, indicating that cell migration was hampered (Figure 2A). Our Transwell assays using PC3, DU145, C4-2, and CWR22r shCCM1 cells revealed that CCM1 suppression reduced both migration and invasion in multiple metastatic PCa cell lines (Figures 2B and S2A–C). Cadherin switching is a phenotype that represents a major hallmark of the metastatic process in most types of epithelial cancer. PC3 shCCM1 cells grown in a 3D hanging drop culture system or 2D culture plate system uniformly exhibited E-cadherin upregulation and N-cadherin downregulation (Figures 2C and 4D, 4F), indicating that CCM1 upregulation can induce E-cadherin and N-cadherin switching. SLUG, SNAIL, and TWIST are representative transcription factors responsible for epithelial-mesenchymal transition (EMT) gene signatures including cadherin switching, which is induced via the repression of E-cadherin and upregulation of N-cadherin (21,22). qPCR analysis indicated that SLUG and TWIST expression was reduced in our shCCM1 cells (Figure 2D), whereas SNAIL expression was not affected (Figure S2E). All primers for qPCR are described in Table 1. Our soft agar assay revealed that CCM1 is necessary for the anchorage-independent survival of PCa cells (Figures 2E and S2D). Suppression of CCM1 prominently reduced the survival of PC3 and DU145 cells in clonogenic survival assays (Figure 2F). Suppression of CCM1 did not result in notable changes in proliferation and cell cycle progression (Figures 2G–H and S2F–G). We conclude that CCM1 increases the migration (motility), invasion, and survival of PCa cells.

**Table 1.**
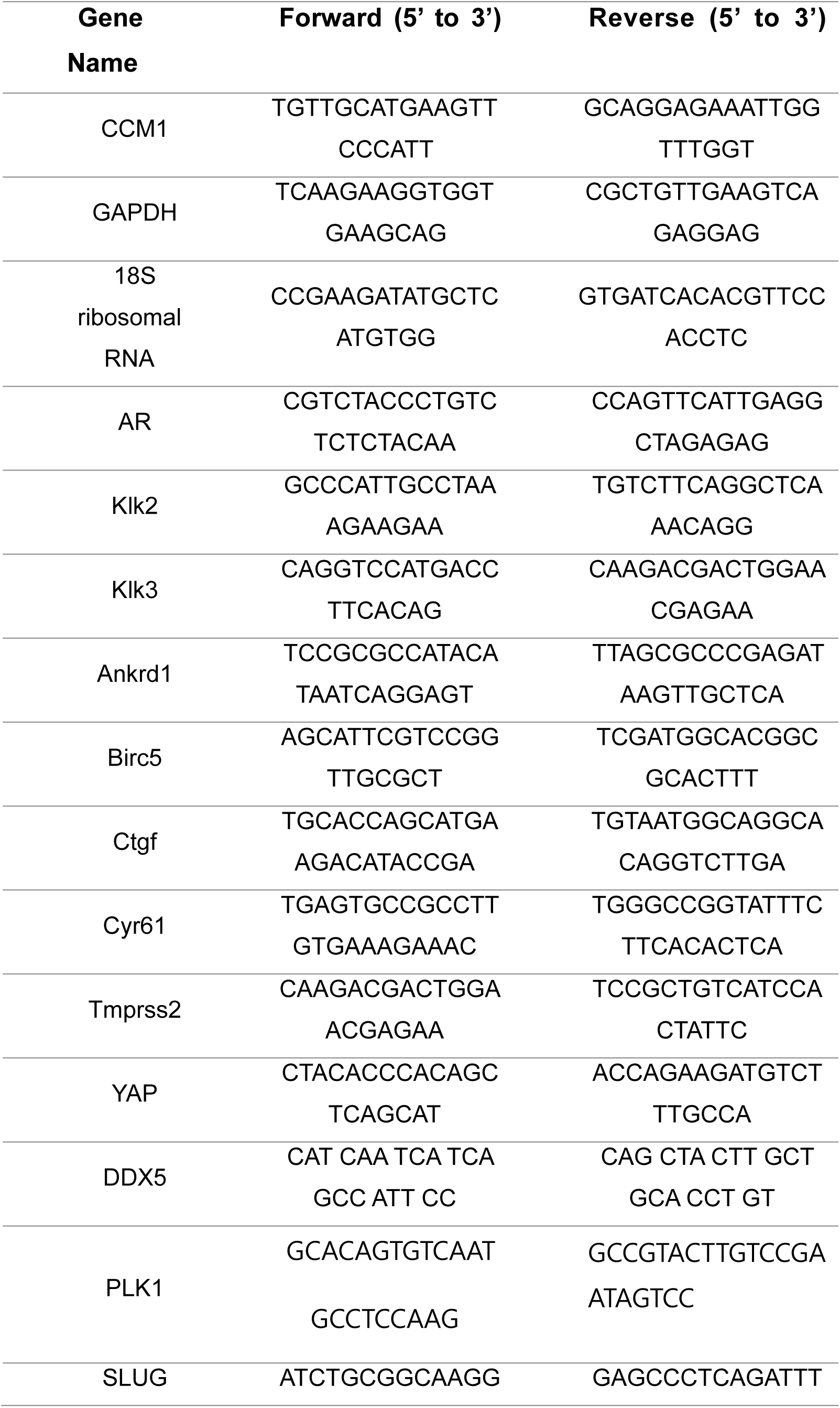
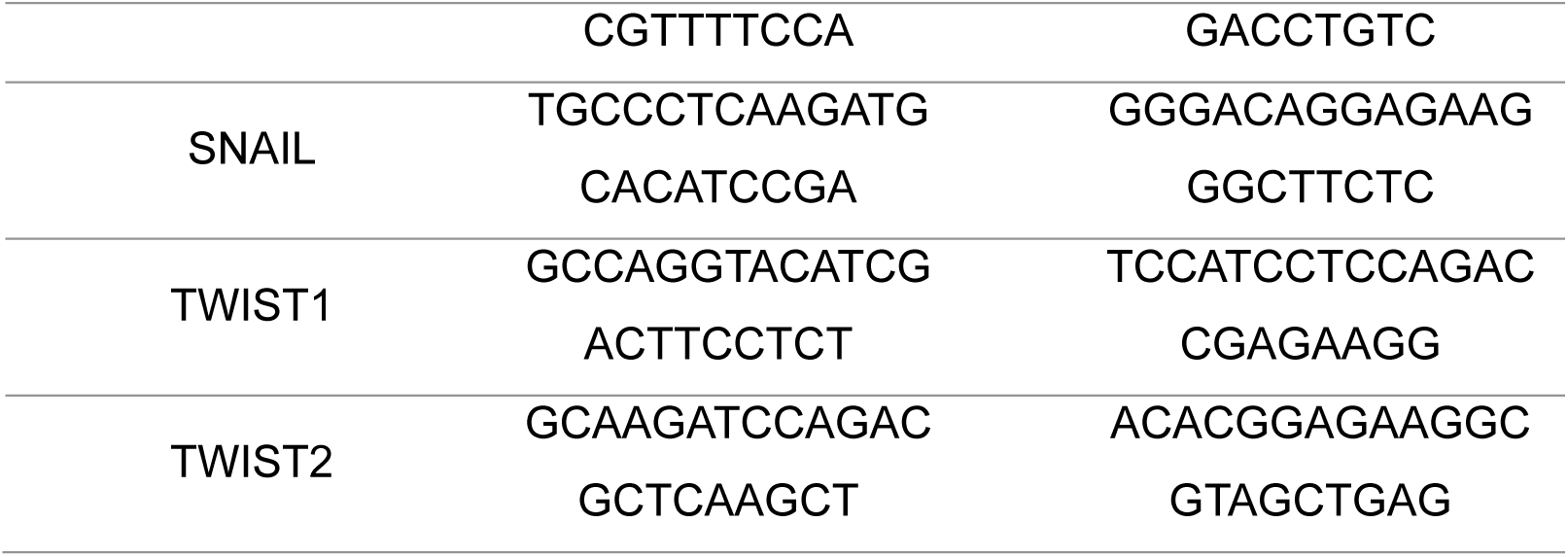
List of qPCR primers used in this study.

**Figure 2.**
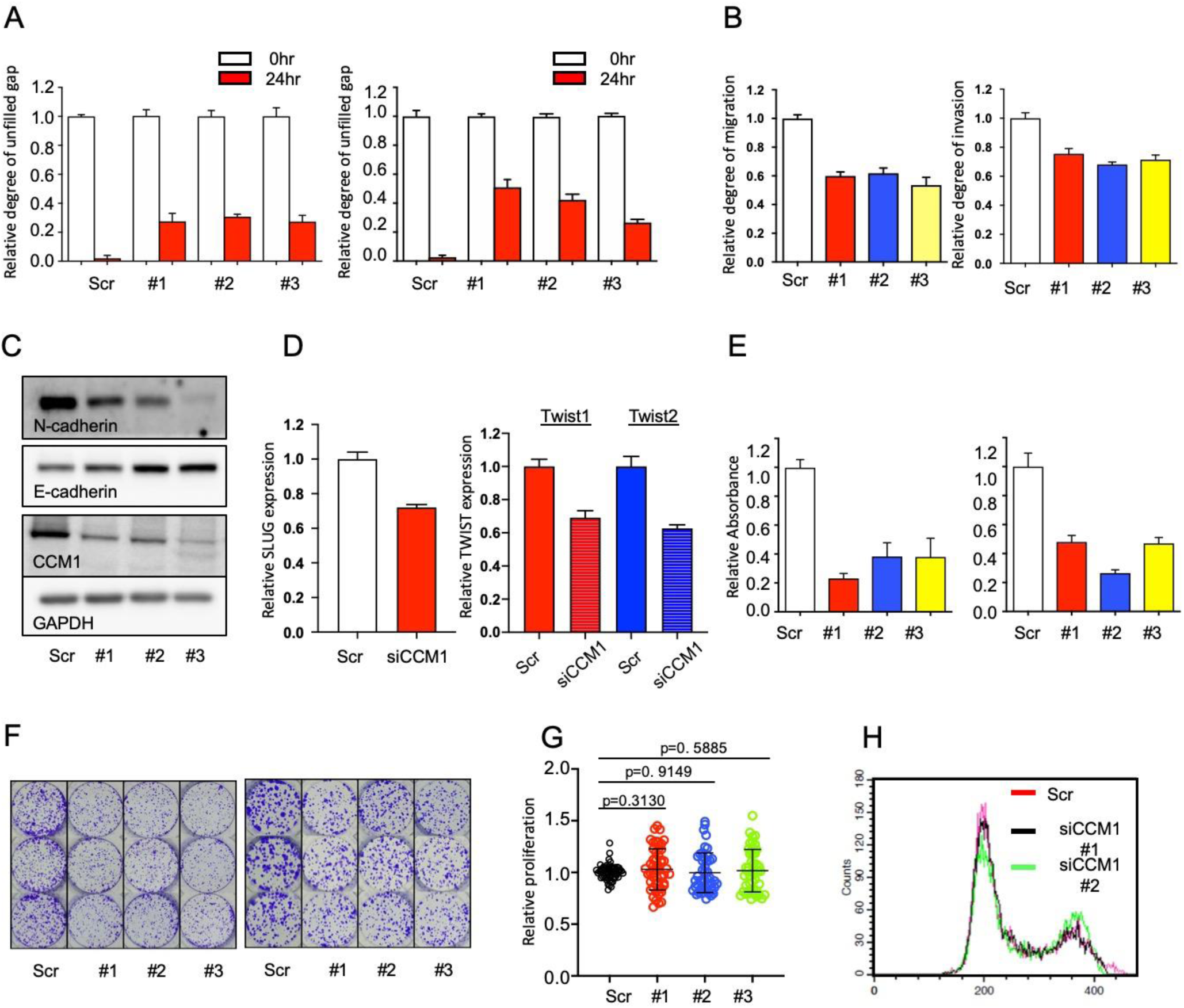
Suppression of CCM1 downregulates metastatic hallmarks in prostate cancer cells *in vitro*. (A) Scratch wounds were created in fully confluent PC3 (left) and DU145 shCCM1 cells (right), and unfilled gaps were measured after 24 h. The data are presented as the mean ± SEM of the unclosed length expressed as a ratio to that at 0 h. (B) PC3 shCCM1 cells were plated on Transwell membranes coated without or with Matrigel for migration (left) and invasion assays (right), respectively, incubated for up to 72 h, and stained. Data are presented as the mean ± SEM of a ratio to that in scramble (Scr) control level. (C) PC3 shCCM1 cells were grown in 3D hanging drop culture for 3 days, and protein extracts were analyzed for changes in N-cadherin and E-cadherin expression via immunoblotting. (D) SLUG expression (left) and TWIST1 and TWIST2 expression (right) were analyzed by qPCR in CCM1-suppressed PC3 and C4-2 cells, respectively. (E) PC3 (left) and Du145 (right) shCCM1 cells were grown in soft agar for 7 days, and lysates were analyzed using a microplate reader to measure anchorage-independent survival. The data are presented as the mean ± SEM of absorbance expressed as a ratio to that in Scr control. (F) PC3 (left) and DU145 shCCM1 cells (right) were plated in each well of a six-well plates, grown for 10 days, and stained with crystal violet for clonogenic survival assays. Representative images from triplicate individual experiments are presented. (G) PC3 shCCM1 cells were plated and grown for 3 days, and cellular proliferation was analyzed using Ez-Cytox assay kits. Scatter plots present the ratio of endpoint values (n = 3–5/set) from five independent experiments relative to the control mean values in each corresponding experiment. Welch’s unpaired *t*-test was performed for statistical analysis. (H) CCM1 was suppressed using two different siRNAs in PC3 cells, and the cell cycle distribution was analyzed. Panels C and H present representative images from three independent experiments, and other graphs in this figure were generated from more than three independent experiments.

### 3. CCM1 regulates YAP/TAZ signaling

As stated previously, accumulated evidence indicates that YAP/TAZ signaling is correlated with PCa metastasis. We thus investigated whether CCM1 upregulates YAP/TAZ signaling. Because aberrant AR signaling is associated with the progression of PCa, we also characterized the effect of AR signaling on CCM1 function. For this purpose, we sorted PC3 cells stably expressing ectopic AR into AR high and AR low clones (Figure S3A). Although WT PC3 cells do not respond to DHT stimulation, they retain multiple AR co-activators and repressors, and the introduction of ectopic AR restores androgen responsiveness in PC3 cells (Figure S3B), as reported previously (23). We also grew multiple types of PCa cells in androgen-depleted, charcoal-stripped FBS to imitate the ADT condition, and dihydrotestosterone (DHT) was administered.

As we expected, overexpression of CCM1 increased the activity of the TEAD reporter (8XGTIIC luciferase) in PC3 and LNCaP cells (Figure 3A). In addition, silencing of CCM1 decreased TEAD reporter activity in LNCaP and C4-2 cells regardless of DHT stimulation (Figure 3B–C). Because CCM1 overexpression modestly increased TEAD reporter activity in PC3, LNCaP (Figure 3A), and C4-2 cells (data not shown), we focused on CCM1 suppression in further studies. Suppression of CCM1 also reduced TEAD activity in DU145 cells (Figure S3C) as well as AR low and AR high PC3 cells regardless of DHT stimulation (Figure S3D), indicating that CCM1-mediated regulation of YAP/TAZ signaling was functional in multiple types of PCa cells. It is known that YAP1 suppression does not completely abolish TEAD reporter activity based on the presence of residual TAZ expression, which we validated (Figure S3E). Using qPCR analysis, we also confirmed that the expression of representative YAP/TAZ target genes such as Ankrd1, Birc5, Ctgf, and Cyr61 was downregulated by the RNAi-mediated silencing of CCM1 (Figure 3D). Suppression of CCM1 by siRNA transfection was effective in multiple types of PCa cells (Figure S3F). Our data indicate that suppression of CCM1 downregulates YAP/TAZ activity regardless of androgen responsiveness in PCa cells and that increased CCM1 expression can upregulate YAP/TAZ signaling.

**Figure 3.**
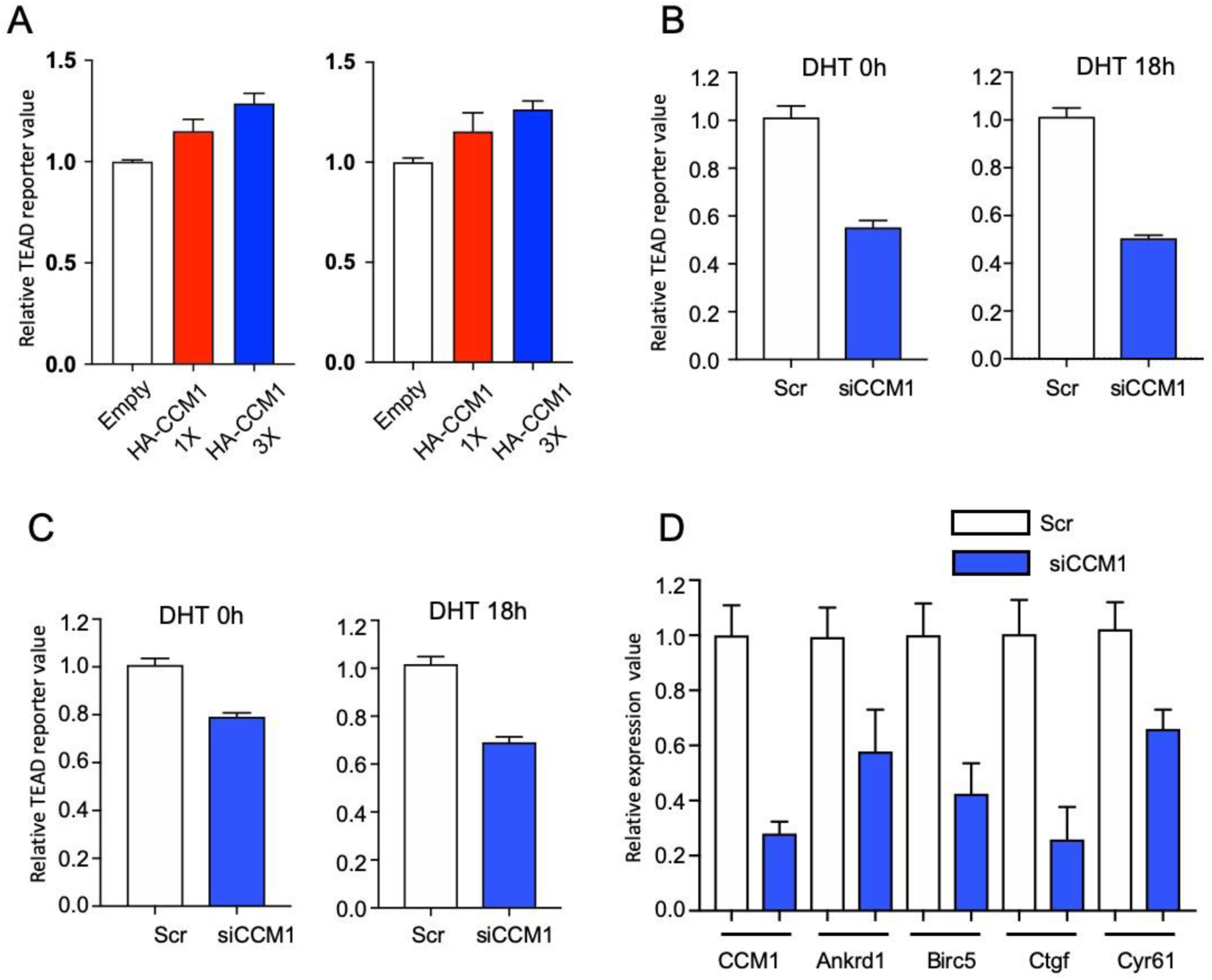
CCM1 upregulates Yes-associated protein (YAP)/transcriptional co-activator with PDZ-binding motif signaling. (A) PC3 (left) and LNCaP (right) cells were co-transfected with increasing amounts of CCM1 overexpression plasmids and the same amounts of TEAD reporter plasmids and grown in normal medium. TEAD reporter activity was measured using luciferase assays. (B) LNCaP and (C) C4-2 cells were co-transfected with CCM1 siRNA and TEAD reporter plasmids. Cells were grown in charcoal-stripped FBS-supplemented medium and stimulated with or without 1 µM DHT for 18 h, as indicated. Reporter activities are shown as relative to that in scramble (Scr) control cells. (D) CCM1 was RNAi-suppressed in LNCaP cells, and the expression levels of CCM1 and representative YAP target genes were analyzed via qPCR. The expression level of each gene is shown relative to that in the Scr control. The data in this figure are presented as the mean ± SEM, and graphs were generated from more than three independent experiments.

### 4. DDX5 is a functional downstream mediator of CCM1 in the regulation of metastatic hallmarks

In our previous proteomics study using CCM1 as a core scaffold protein to identify phosphorylation sites in the CCM complex (24), we identified DDX5 as a novel CCM1-interacting protein (Table S1). DDX5, also named p68, is a member of the DEAD box family of RNA helicases that contain nine conserved motifs, including the conserved Asp-Glu-Ala-Asp (DEAD) motif. DDX5 is aberrantly expressed/modified in several types of cancers, suggesting that it plays important roles in cancer development and progression (25-27). We validated the interaction between endogenous DDX5 and Flag-tagged CCM1 in co-immunoprecipitation (co-IP) and reverse co-IP assays (Figure 4A–B). Our co-IP assay also demonstrated that DDX5 was immunoprecipitated with WT CCM2 (Figure S4A). Because CCM2 is a major interacting protein of CCM1 (28) and the CCM1-CCM2 interaction was confirmed from our PC3 prostate cancer cell lysates (Figure S4B), we further investigated whether CCM1 is required for the CCM2-DDX5 interaction. Compared with the results of WT CCM2 overexpression, neither IP of L198R CCM2 mutant (CCM1 nonbinding) nor suppression of CCM1 resulted in a notable decrease in the amount of co-immunoprecipitated HA-DDX5 (Figures 4C, S4C–D). Our data indicate that CCM1 was not required for the CCM2-DDX5 interaction.

**Figure 4.**
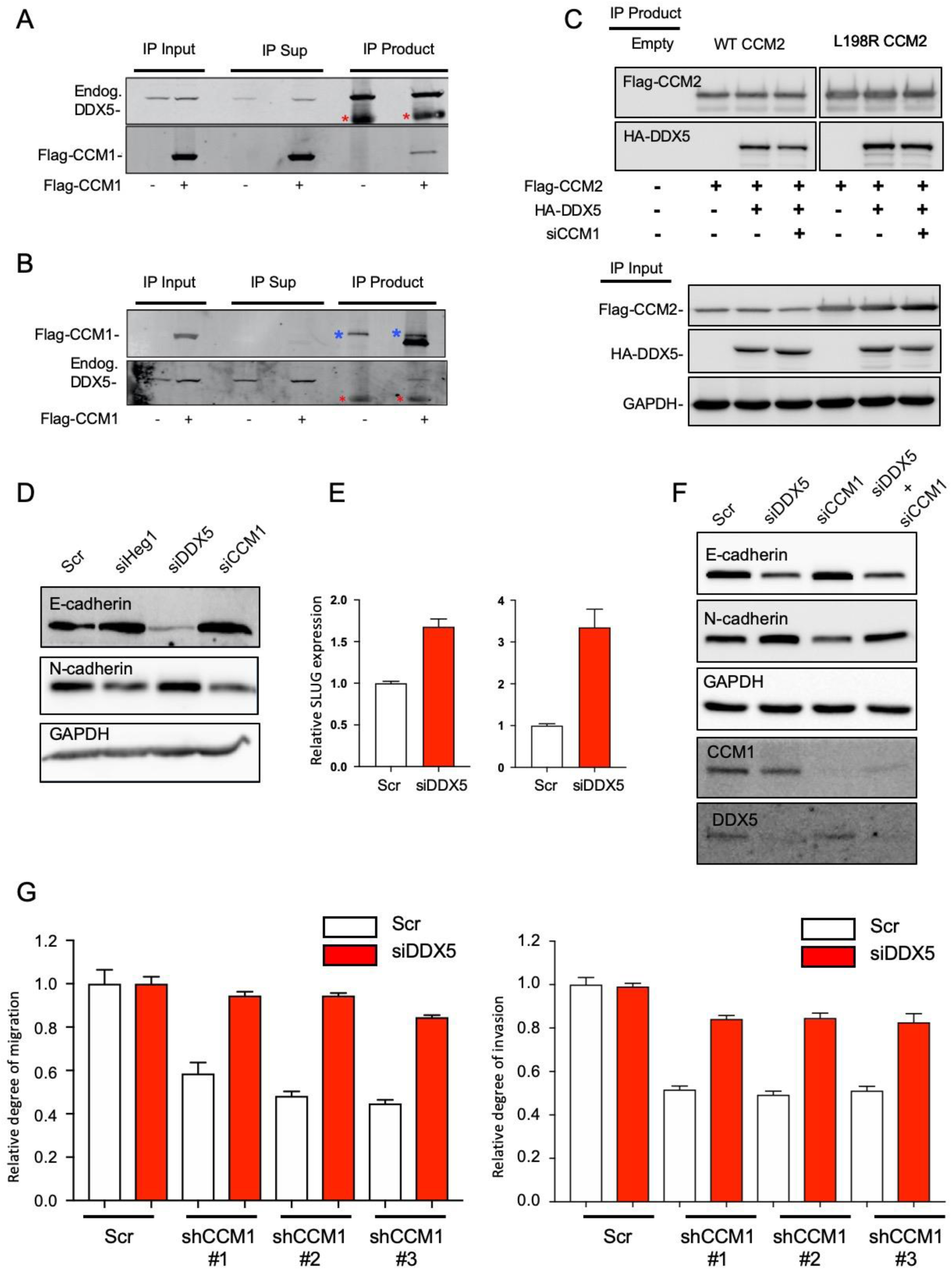
DDX5 interacts with the CCM complex and functions as a mediator of CCM1. (A) CCM1 was co-immunoprecipitated using a DDX5 immunoprecipitation assay, and (B) DDX5 was co-immunoprecipitated in a CCM1 immunoprecipitation assay using U2OS cell lysates. Red asterisks: heavy chain, blue asterisks: non-specific. (C) HA-tagged DDX5 overexpression plasmid or empty plasmid was transfected into PC3 cells stably overexpressing Flag-tagged WT CCM2 or L198R CCM2. Additionally, CCM1 was RNAi-silenced in WT or L198R CCM2-expressing cells transfected with HA-DDX5 overexpression plasmids. (D) Suppression of HEG1, DDX5, and CCM1 using siRNA was performed in PC3 cells, and changes in E-cadherin and N-cadherin expression were analyzed. (E) DDX5 was suppressed in PC3 (left) and DU145 (right) cells, and SLUG expression was analyzed using qPCR. (F) Individual RNAi silencing of DDX5 or CCM1 and co-silencing of DDX5 and CCM1 were performed in PC3 cells to compare changes in E-cadherin and N-cadherin levels. (G) DDX5 expression was suppressed in C4-2 shCCM1 cells, which were plated on Transwell membranes coated without or with Matrigel for migration (left) and invasion assays (right), respectively, incubated for up to 72 h, and stained. The data are presented as the mean ± SEM of a ratio to DDX5 expression in intact scramble control cells. Representative images were presented from three independent experiments, and graphs were generated from three independent experiments.

Because suppression of CCM1 reduced several metastatic hallmarks, we further investigated cadherin switching following RNAi-mediated silencing of CCM1 or CCM1-interacting proteins. Suppression of CCM1 and HEG1 downregulated N-cadherin and upregulated E-cadherin expression (Figure 2C, 4D). HEG1 is a major CCM1-interacting protein with uncertain functions (29), and CCM and HEG1 were also found to interact genetically (30). Contrary to the changes in cadherin expression after the suppression of CCM1 or HEG1, suppression of DDX5 clearly upregulated N-cadherin and downregulated E-cadherin expression (Figure 4D). Our data indicate that suppression of CCM1 and HEG1 inhibited cadherin switching, whereas suppression of DDX5 induced cadherin switching. Accordingly, suppression of DDX5 upregulated SLUG (Figure 4E), which was contrary to our finding that suppression of CCM1 led to SLUG downregulation (Figure 2D). To identify whether DDX5 is a functional downstream mediator of CCM1, we used a co-RNAi silencing strategy for DDX5 and CCM1. Co-suppression of DDX5 and CCM1 decreased E-cadherin and increased N-cadherin expression similarly as observed for silencing of DDX5 alone, restoring the suppression of CCM1-induced changes in cadherin levels (Figure 4F). Next, we suppressed DDX5 in three independent C4-2 shCCM1 cell lines and confirmed that co-suppression of DDX5 restored the migration and invasion of CCM1-silenced PCa cells (Figure 4G). Our data indicate that DDX5 is a major functional downstream mediator of CCM1 in the regulation of metastatic hallmarks such as the migration and invasion of PCa cells.

To reveal the potential impact of genomic rearrangements in the regulation of CCM1 and DDX5 genes, we explored 968 whole-genome sequences of PCa downloaded from four different cohorts including the Canadian prostate cancer genome network (31-33). Then, we compared the frequency of structural variations in the vicinity of these genes, including copy number alterations and genomic rearrangements. Although both the CCM1 and DDX5 gene loci (7q21.2 and 17q23.3, respectively) are often amplified in a substantial fraction of PCa tumors (approximately 10% for CCM1 and approximately 5% for DDX5), these rates were not increased in metastatic cancer tissues (Figure S5A). In addition, we did not find any rearrangement candidates near either gene (data not shown). Although it is believed that DDX5 is upregulated in several tumors compared with its levels in matched normal tissues (34), our PCTA and independent Oncomine analyses of two PCa datasets (35,36) revealed the downregulation of DDX5 expression in metastatic PCa compared with its expression in primary PCa (Figure S5B–C).

### 5. DDX5 suppresses YAP/TAZ signaling through PLK1

To investigate whether DDX5 regulates YAP/TAZ signaling, we suppressed DDX5 in multiple types of PCa cells. Contrary to the effects of CCM1 suppression, suppression of DDX5 increased TEAD reporter activity in LNCaP, C4-2, and PC3 AR low and high cells (Figure 5A–C). Accordingly, overexpression of WT DDX5 suppressed TEAD reporter activity in all PCa cell types tested (Figures 5D–E and S6C). Our data indicate that DDX5 can uniformly downregulate YAP/TAZ signaling in multiple types of PCa cells regardless of their androgen responsiveness. PC3 AR high cells exhibited higher TEAD reporter activity than PC3 AR low cells (Figure 5C). It has been reported that the phosphorylation of DDX5 at Y593 is important for EMT in HT29 colon cancer cells (27). To characterize whether Y593 phosphorylation is required for the DDX5-mediated regulation of YAP/TAZ signaling, we overexpressed WT DDX5 or non-phosphorylatable Y593F DDX5 in LNCaP, C4-2, C4-2B, and CWR22r cells. Overexpression of WT DDX5 uniformly suppressed YAP/TAZ activation in multiple PCa cell lines, whereas overexpression of Y593F DDX5 did not alter YAP/TAZ activation (Figures 5D–E and S6C). Increasing ectopic WT DDX5 levels further suppressed TEAD reporter activity, whereas overexpression of Y593F DDX5 had no such effect (Figure 5E), indicating that Y593-phosphorylated DDX5 was required for the downregulation of YAP/TAZ signaling. Co-suppression of DDX5 and YAP1 downregulated TEAD reporter activity to a similar level as suppression of YAP1 alone (Figures 5F and S6A). Our data indicate that DDX5 regulates YAP/TAZ signaling through YAP rather than TAZ co-activator.

**Figure 5.**
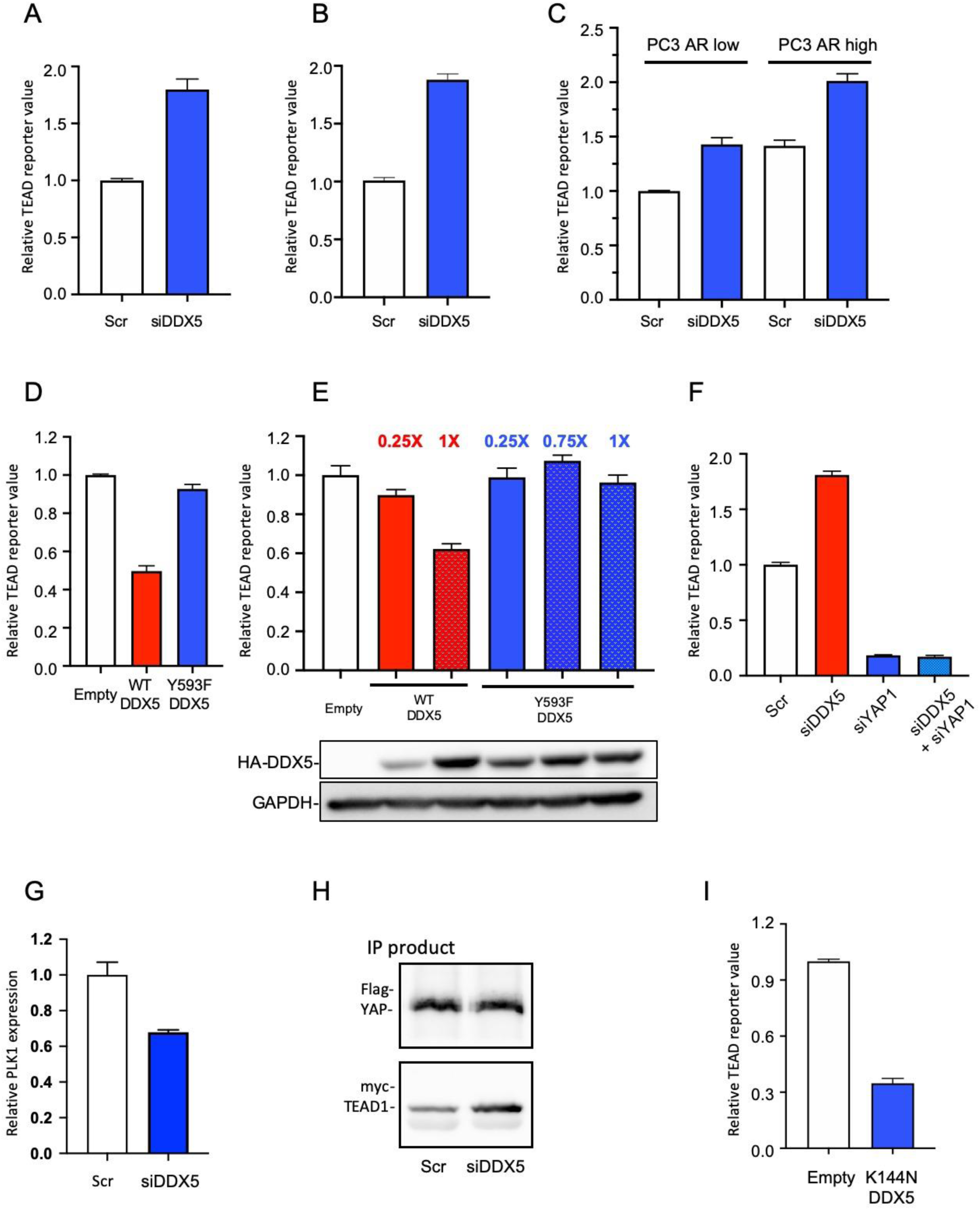
DDX5 suppresses Yes-associated protein (YAP)/transcriptional co-activator with PDZ-binding motif signaling in prostate cancer cells. DDX5 was RNAi-suppressed in (A) LNCaP, (B) C4-2, and (C) PC3 cells with low or high androgen receptor (AR) expression, grown in regular medium, and TEAD reporter activities were analyzed. (D) WT DDX5 or Y593F DDX5 was overexpressed in C4-2 cells, and lysate was analyzed using the TEAD reporter assay. (E) WT or Y593F DDX5 was overexpressed in C4-2 cells, and each lysate was analyzed using the TEAD reporter assay. The representative expression levels of HA-DDX5 are shown below the graph. (F) RNAi suppression of DDX5 or YAP1 alone or co-suppression of DDX5 and YAP1 was performed, and TEAD reporter activity was analyzed in LNCaP cells grown in charcoal-stripped FBS and stimulated with DHT for 18 h. (G) DDX5 was suppressed in PC3 cells, and the mRNA expression level of PLK1 was analyzed via qPCR. (H) Flag-tagged YAP was immunoprecipitated from the nuclear lysates of PC3 cells co-transfected with Flag-tagged YAP, myc-TEAD1 overexpression plasmids, and siDDX5. The resultant immunoprecipitation products were immunoblotted with anti-Flag and anti-myc antibodies. (I) K114N helicase-dead mutant DDX5 was overexpressed in LNCaP cells, and TEAD reporter activity was analyzed. The data are presented as the mean ± SEM relative to the expression in scramble control or empty plasmid-transfected control cells. Graphs were generated from three independent experiments. Empty: empty plasmid transfection.

Previous research found that YAP does not interact with DDX5 in HaCaT cells (37), and we also observed using a co-IP assay that DDX5 did not interact with YAP1 in PCa cells (data not shown), precluding the possibility that DDX5 regulated YAP/TAZ signaling through its interaction with YAP. Because it has been reported that DDX5 upregulates PLK1 (38) and PLK1 downregulates YAP/TAZ signaling by disrupting the interaction between YAP and TEAD (39), we investigated whether DDX5 regulated YAP/TAZ signaling through PLK1 in PCa cells. We found that PLK1 expression was reduced upon DDX5 suppression (Figure 5G), and our co-IP experiments using PC3 nuclear lysates revealed that the YAP-TEAD interaction was increased upon DDX5 suppression (Figures 5H and S6B). Our data indicate that PLK1 was responsible for the regulation of YAP/TAZ signaling by DDX5 in PCa cells. DEAD box RNA helicases use RNA as a substrate to stimulate ATPase activity, and the energy is then used to displace duplex RNA or proteins that bind the RNA substrate (40). Overexpression of helicase-dead K144N mutant DDX5 also decreased TEAD reporter activity with similar efficiency as WT DDX5, indicating that the RNA helicase activity of DDX5 was not involved in the regulation of YAP/TAZ signaling (Figure 5I).

Additionally, we also investigated changes in major mechanisms regulating YAP/TAZ signaling such as alterations in the expression or subcellular localization of YAP following DDX5 or CCM1 suppression. The subcellular localization of YAP was not changed upon DDX5 suppression (Figure S6D). Both the expression and subcellular localization of YAP were not changed by CCM1 suppression (Figure S6E–F). Although our finding that DDX5 interacts with TEAD1 suggests that DDX5 may be a novel co-repressor of YAP/TAZ signaling, Y593 phosphorylation of DDX5 did not alter the strength of this interaction (Figure S7A–B). Overexpression of WT or Y593F DDX5 did not change the expression of total YAP1 (Figure S7C) or the subcellular localization of total YAP1 and S127-phosphorylated YAP (Figure S7D) compared with the effects of empty plasmid transfection.

As stated previously, YAP/TAZ signaling is a positive regulator of AR signaling, and it is suggested to be responsible for CR in prostate cancer (12,13). We found that the suppression of DDX5 and YAP1 prominently upregulated and downregulated ARR reporter activity, respectively (Figure S7E–F). Our data indicate that DDX5 can indirectly upregulate AR signaling via its regulation of YAP/TAZ signaling and that this activity is mediated through YAP rather than TAZ. Accordingly, we also found that suppression of CCM1 downregulated ARR reporter activity in LNCaP (Figure S7G), CWR22r, C4-2, and C4-2B cells (data not shown). Suppression of CCM1 also downregulated representative AR target genes (Figure S7H). Our data indicate the existence of a functional CCM1-DDX5-PLK1-YAP-AR signaling pathway in PCa cells.

### 6. CCM1 regulates the phosphorylation of DDX5 at Y593

Because phosphorylation of DDX5 at Y593 was necessary for the suppression of YAP/TAZ signaling and CCM1 and DDX5 had opposite regulatory effects on YAP/TAZ signaling, we hypothesized that CCM1 may downregulate the function of DDX5 by blocking its phosphorylation at Y593. CCM1 was silenced in PC3 cells and non-prostate cancer cell lines (U2OS and HT-29), followed by stimulation with WNT3a (Figure 6A–B) or PDGF ligand (Figure 6B). In addition to PDGF, which was previously revealed to induce the phosphorylation of DDX5 at Y593 (27), we found that WNT stimulation also induced phosphorylation at Y593. Notably, suppression of CCM1 further upregulated the basal level of Y593-phosphorylated DDX5 in all cell lines grown in serum-free medium, similarly as observed in scramble control cells stimulated with WNT or PDGF (Figure 6A–B), indicating that the CCM1-mediated regulation of Y593 phosphorylation was not limited to PCa cells. As observed for CCM1 suppression, suppression of HEG1 also increased Y593 phosphorylation in DDX5 (Figure 6A). Notably, suppression of HEG1 and CCM1 produced an identical pattern of cadherin switching (Figure 4.D)

**Figure 6.**
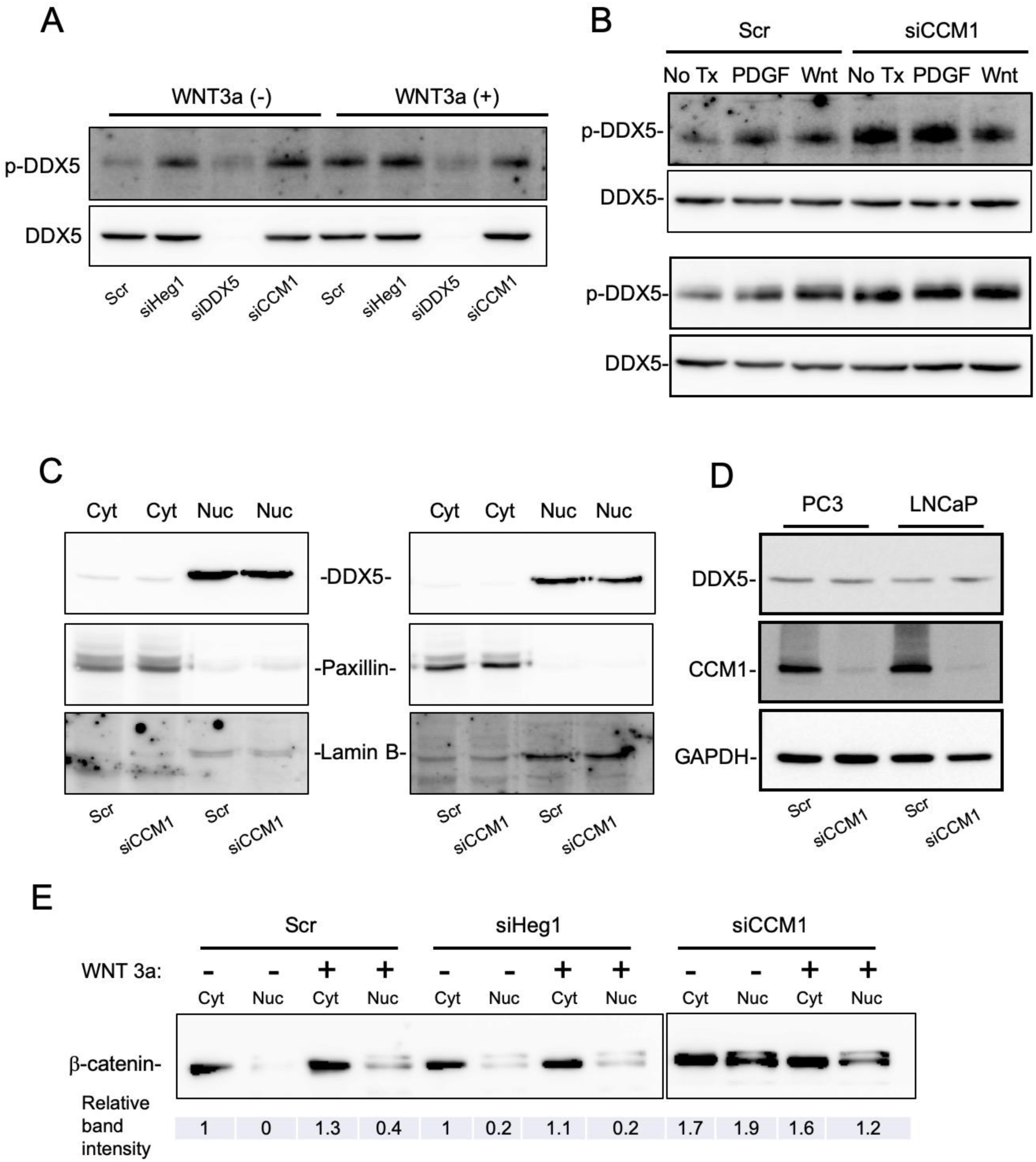
CCM1 regulates the phosphorylation of DDX5 at Y593. (A) PC3 cells were RNAi-silenced as indicated and stimulated with WNT3a. (B) CCM1 was silenced in U2OS (top) and HT29 (bottom) cells using RNAi, followed by stimulation with WNT3a or PDGF ligand. (C) CCM1 was RNAi-silenced in PC3 (left) and LNCaP (right) cells, and they subcellular localization of DDX5 was analyzed. Cytosolic and nuclear fractions were subjected to immunoblotting with antibodies against paxillin and lamin B to target the cytosolic and nuclear fractions, respectively. (D) DDX5 and CCM1 expression was analyzed in CCM1-silenced PC3 and LNCaP cells. (E) The subcellular localization of β-catenin in CCM1-silenced PC3 cells treated with/without Wnt3a ligand. Relative band intensity acquired via densitometry is shown below the figure. p-DDX5: Y593-phosphorylated DDX5, Cyt: cytosolic fraction, Nuc: nuclear fraction.

Next, we investigated whether CCM1 regulates the subcellular localization or expression of DDX5. Our subcellular fractionation experiment demonstrated that the RNAi-mediated silencing of CCM1 did not change the subcellular localization of endogenous DDX5 (Figure 6C) or its expression level (Figure 6D). Because the suppression of CCM1 and WNT3a ligand upregulates WNT signaling and induces the subsequent nuclear import of β-catenin (41), the subcellular localization of β-catenin was evaluated as a control for our subcellular fractionation study (Figure 6E). Suppression of HEG1, in contrast to the effects of CCM1 suppression, did not induce the nuclear mobilization of β-catenin.

In conclusion, our data indicate that suppression of phosphorylation at Y539 was the mechanism by which CCM1 downregulated the function of DDX5 and that CCM1 upregulated YAP/TAZ signaling through two successive processes, namely the negative regulation of DDX5 by CCM1 and the negative regulation of YAP/TAZ signaling by DDX5 through PLK1.

## Discussion

Many recent findings demonstrated that YAP/TAZ and AR signaling serve as the central regulatory mechanisms for prostate tumorigenesis and several cancer-associated pathways can promote YAP/TAZ signaling (14,42). However, the complete mechanism by which YAP/TAZ signaling becomes hyperactivated and interacts with the stroma, as well as their precise role in PCa development, has not been elucidated. We propose that CCM1 is an important regulator of the YAP/TAZ and AR signaling in PCa cells. That stated, our evidence indicates that despite the concomitant activation of multiple pro-tumoral signaling pathways, suppression of CCM1 ameliorated representative metastatic hallmarks, namely motility, invasion and survival (43), in multiple types of both androgen-responsive and androgen–non-responsive metastatic PCa cells. CCM1 may not be sufficient to induce tumorigenesis, but it appears to unleash the oncogenic potential of multiple cellular signals, promoting cancer progression including the invasiveness and survival of PCa cells. Suppression of DDX5 or CCM1 changed the expression levels of Slug and Twist, but not Snail, although all three are generally considered important transcription factors involved in EMT in cancer cells. It was demonstrated that the negative feedback between SLUG and SNAIL is differentially regulated in highly and lowly invasive cancer cells (21). In breast cancer, SNAIL is involved in early EMT, and decreased SNAIL and increased SLUG expression upregulate phospholipase 2, which is correlated with the increased invasiveness in cancer cells. This may explain why SLUG, but not SNAIL, was notably affected in highly metastatic PCa cells.

Based on genetic aberrations of AR or the increased synthesis of androgens (in the tumor microenvironment [TME]) or adrenal androgen precursors, it has been reported that AR expression is increased (44) and that AR signaling is consistent with the castration level of androgens in patients with mCRPC. However, whether AR signaling is physiologically or supra-physiologically potentiated in the TME of patients with mCRPC is an important open question (45,46). Our data suggest that elevated levels of CCM1 can potentiate both ligand-dependent and ligand-independent AR signaling in the TME through the upregulation of YAP/TAZ signaling and that the resultant increase in AR signaling may also be important for progression to advanced prostate cancer. Approximately 15%–25% of patients with CRPC do not respond to first-line treatment with the “supra-castration” agents abiraterone and enzalutamide (47), and it would be interesting to investigate whether CCM1 is responsible for the non-responsiveness.

Most research on CCM has been conducted in the field of vascular biology, and there is limited understanding regarding its function in cancer biology (41,48). Because of the multifocal and heterogeneous nature of human PCa, in which distinct molecular and genetic alterations are associated with various “clones,” the unique and prominent increase of CCM1 expression only in mCRPC was an unexpected finding. Several previous studies investigated genomic alterations in primary PCa and mCRPC and revealed important oncogenic changes responsible for the progression of PCa (3,49). However, until recently, few large-scale sequencing studies of men with metastatic lethal PCa have been conducted (50). In the era of PSA screening, most PCa lesions are well differentiated and clinically localized at diagnosis with extremely few lethal events, and these cohorts largely consist of cases that are understood to be de-enriched in men carrying significant genetic risk factors for aggressive disease. In addition, the identification of various somatic mutations responsible for treatment resistance, including CR, from genomic studies of gene mutations and chromosomal DNA arrangements alone without transcriptomic data from heavily treated cancer samples may not reveal selective and crucial alterations in cellular signaling events. These facts may explain why CCM1 has not been previously identified in PCa biology. As the scope of large-scale genomic and transcriptomic analysis of metastatic cancers is mostly confined to heavily treated mCRPC (51), we were unable to differentiate non-treated pure metastatic events from the pool of all metastatic events in heavily treated CRPCs in our PCTA data. Therefore, whether ADT is responsible for the induction of CCM1 and whether CCM1 is responsible for CR, in addition to uncontrollable metastasis in patients, will remain outstanding questions for further research.

Our data indicate that DDX5 downregulated YAP/TAZ signaling by blocking the PLK1-mediated suppression of the YAP-TEAD interaction, suggesting that DDX5 may also regulate other nuclear events such as the recruitment of YAP-associated activators or repressors to their target gene promoters. Although DDX5 is known to promote proliferation and tumorigenesis in other cancers in addition to PCa, DDX5 downregulation also activates a pro-survival pathway involving mTOR and MDM2 signaling, leading to the inhibition of pro-apoptotic activity in HeLa cells (52), revealing the pleiotropic function of DDX5 depending on the cellular context. For example, contrary to our findings, phosphorylation of DDX5 at Y593 induced by c-Abl was reported to induce EMT in colon cancer cells (27).

It is believed that DDX5 expression is upregulated in multiple cancer types. Contrarily, we observed that DDX5 expression was downregulated in metastatic PCa compared with that in primary PCa, although we found no evidence of genomic rearrangement. These findings, at least in part, indicate that the previously reported function of DDX5 in other cancer models may not be directly applicable to the biology of PCa. Of note, our functional analysis consistently revealed that DDX5 was a downstream mediator of CCM1 and that DDX5 suppressed a metastatic hallmark in multiple types of PCa cells, adding another layer of complexity to understanding the function of DDX5. Of note, considering the suppressive effects of DDX5 on YAP/TAZ signaling, downregulation of DDX5 expression in patients with mCRPC appears to further potentiate the CCM1-mediated upregulation of YAP/TAZ signaling and metastasis. Because we never observed a notable change in DDX5 expression upon the RNAi-induced suppression of CCM1 in PCa cells (data not shown) or the genomic rearrangement rates of DDX5 gene between primary PCa and mCRPC, these findings suggest that mechanisms other than CCM1 signaling, such as unidentified epigenetic changes, were responsible for the downregulation of DDX5 expression in the mCRPC samples. In addition to DDX5, we also attempted to identify the cause of CCM1 induction but found that gene rearrangement was not responsible. Therefore, our data indicate that other molecular mechanisms, such as epigenetic changes or cues from the TME, may upregulate CCM1 in metastatic tissues.

Importantly, the incidence of PCa metastasis has dramatically increased because of advancements in therapeutic strategies for patients, indicating the importance of preventing these detrimental metastatic events. We uncovered an interesting scenario in which PCa cells arising in disparate genetic backgrounds appear to share common central molecular events regarding progression. To conclude, we are convinced that CCM1 bears tremendous potential for molecular oncology, and we await future discoveries regarding its function, regulation, and potential diagnostic and therapeutic manipulations at the bedside.

## ACKNOWLEDGMENTS

This work was supported by the Basic Science Research Program, through the National Research Foundation of Korea (NRF), funded by the Ministry of Science, ICT & Future Planning (NRF-2017R1D1A1B03029063). We thank professor Mark H. Ginsberg and Jong-Sup Bae for carefully reading and providing helpful comments on this manuscript.

## Methods

### 1. Cell culture and reagents

Human DU145, PC3, LNCaP, C4-2, and CWR22r cells were grown in RPMI medium (Welgene) supplemented with 10% FBS (Welgene) and 1% penicillin/streptomycin (Invitrogen) at a temperature of 37°C in an atmosphere of 5% CO_2_. To induce androgen deprivation, RPMI medium supplemented with 10% charcoal-stripped FBS (Scipak) was used. We used Ambion™ Silencer™ Select Validated siRNAs for gene silencing of CCM1 and DDX5.

### 2. Generation of stable cell lines

To generate lentiviruses for silencing CCM1, Lenti293 packing cells were plated at 6 × 10^5^ cells/well in six-well tissue culture plates and transfected with pLKO.1-puro lentiviral empty vector or shCCM1 vector (Sigma) using X-treme GENE HP DNA transfection Reagent (Roche). Transfected 293 cells were grown in DMEM containing 30% FBS for 24 h or 48 h. The conditioned medium containing lentiviruses was collected and pooled. Hexadimethrine bromide (Sigma) was added at a final concentration of 8 µg/ml in cell culture medium, and infection was performed by adding lentiviral conditioned medium to PCa cells overnight. Starting the following day, cells were treated with puromycin for 7 days to select lentivirus-infected cells.

### 3. Immunoblotting assay

Cells with stably CCM depletion (shCCM1 cells) via lentiviral shRNA particle transfection were grown to 80% confluence in six-well or 100-mm culture plates. Then, the cells were lysed in RIPA buffer (50 mM Tris–HCl [pH 7.4], 150 mM NaCl, 1% NP-40, 1 mM EDTA, 1 mM EGTA, 2 mM Na_3_VO_4_, 2 mM NaF, 0.25% sodium deoxycholate) containing a cocktail of protease inhibitors (Roche). The total protein level was quantified using a BCA protein assay kit (Thermo) according to the manufacturer’s protocol. In total, 30–40 µg of total protein were loading into each well of a 10% SDS-PAGE gel and separated using an electroporation system (Bio-Rad). Proteins were transferred onto an NC membrane (Whatman), which was blocked for 1 h with 5% skim milk in Tris-buffered saline containing Tween-20 (TTBS) at room temperature. The membranes were later incubated with primary antibody against CCM1 (Abcam), N-cadherin (BD), E-cadherin (Santa Cruz), DDX5 (Abcam) and GAPDH (Protein Tech) in 5% BSA-TTBS overnight at 4°C and then HRP-conjugated goat anti-rabbit IgG (Jackson) or goat anti-mouse IgG(Sigma) secondary antibodies for 1 h at room temperature. The blots were developed using ECL detection reagent (ATTO).

### 4. Real-time PCR (qPCR) analysis

Total RNA from cells was extracted using TRIzol reagent (TAKARA), and RNA levels were quantified using spectrophotometry. For each reaction, 1 µg of total RNA served as a template for cDNA synthesis with a PrimeScript™ 1st strand cDNA Synthesis Kit (Takara). GAPDH and 18S ribosomal RNA were amplified as reference genes for target mRNA. STBR® Primix EX Taq™ II (Takara, Madison, WI, USA) was employed for qPCR analysis.

### 5. Wound-healing assay

For monolayer wound-healing assays, cells were plated to confluence in 12-well tissue culture plates. Cell layers were scratched with a 200-µl pipette tip and gently washed with PBS twice, and culture medium was gently added to not perturb the cell monolayer. Migration of cells across the wound edge was observed and photographed at 24 h after wounding.

### 6. Transwell invasion and migration assay

Cell invasion and migration were analyzed using 24-well plate format transfer chambers (SPL) with 8-µm sized pores, and we used Matrigel (Corning) to coat membranes in the transfer chambers for invasion assays. Cells (5 × 10^5^ cells/well) were seeded in each transfer chamber. Medium containing 2% FBS was added to each well of a 24-well plate. The Matrigel invasion chambers were transferred to wells containing the chemoattractant using sterile forceps. A suspension of cells in serum-free medium was loaded into the chambers. Cells were incubated for 24–72 h. The invading cells were fixed with 10% formalin for 15 min at room temperature and stained with hematoxylin for 30 min, and the non-invading cells were removed via scrubbing with a cotton tipped swab. The inserts were washed, and the membrane was photographed using a microscope at ×20 magnification.

### 7. Soft agar assay

To assess colony formation, the CytoSelect 96-Well Cell Transformation assay (Cell Biolabs) was used according to the manufacturer’s instruction. Briefly, PCa cells were seeded in soft agar at 2500 cells/well. After 7 days of incubation at 37°C in 5% CO_2_, colony formation was quantified by solubilizing soft agar, lysing cells, and incubating cell lysates with the CyQUANT GR Dye (Cell Biolabs) followed by analysis.

### 8. Hanging drop culture

Cells were counted using an automated cell counter (EVE), and the cell count was adjusted to 1 × 10^6^ cells/ml. Then, 5 ml of PBS were added to the bottom of the dish to preserve humidity for cell culture. A 20-µl pipette was used to deposit 20-µl drops onto the bottom of the lid. The tissue culture plate lid was inverted, placed in the PBS-filled bottom chamber, and incubated at 37°C and 5% CO_2_. Hanging drops were monitored daily for 3 days. After 3 days, each drop was pooled and transferred to a 1.5-ml small tube. Centrifugation was performed at 1200 rpm for 3 min at 4°C, and the resulting pellet was extracted using RIPA buffer for further studies.

### 9. Immunoprecipitation

Transfected cells were allowed to reach a concentration of 1 µg/µl, and 100 µl were left as the input. In total, 40 µl of Protein G-agarose bead suspension (Santa Cruz) were added to 500 µg of lysates and incubated for 1.5 h at 4°C. After centrifugation, the supernatant was transferred to the new tube. One microgram of the primary antibody was added to the cleared lysates, which were inverted and incubated for 1 h at 4°C. Protein G-agarose beads were added to the lysates, which were inverted and incubated overnight at 4°C. Beads were washed three times with RIPA buffer at 4°C before bound proteins were eluted with 2× SDS sample buffer and loaded onto SDS-PAGE gels.

### 10. Reporter assay

In total, 8 × 10^4^ cells were plated in each well of a six-well tissue culture plate 16 h before transfection, which was performed using X-treme GENE HP DNA transfection Reagent (Roche) with specific reporter plasmids. A β-galactosidase plasmid was co-transfected to normalize transfection efficiency. After 7 h, the cells were transferred to a new six-well tissue culture plate and incubated for 2 days with medium containing 10% FBS. Subsequently, the cells were washed with PBS and lysed with 200 µl of reporter lysis buffer. Then, luciferase reporter activities were assayed as relative light units using a luminometer. The experiments were performed in triplicate, and the results were reported as the mean ± SD. To analyze YAP/TAZ activation, a TEAD reporter (8XGTIIC-luciferase) plasmid was used (Addgene). For AR signaling, an ARR2-TK reporter plasmid was kindly provided by Dr. Robert Matusik.

### 11. Clonogenic cell survival assay

Cells from a stock culture were plated following the plating of PC3 or DU145 cells into six-well cell culture plates at 1000 cells/well. After 10 days of incubation at 37°C in 5% CO_2_, medium was removed, and cells were washed with PBS twice. Fixation and staining of cones were performed using 0.4% crystal violet for 30 min. After removing the staining solution, cells were washed with water twice and dried at room temperature.

### 12. Molecular cohort analysis

Our PCTA includes more than 4600 clinical PCa specimens. Based on an extensive survey of public resources, we collected 50 PCa datasets from three public databases: Gene Expression Omnibus (http://www.ncbi.nlm.nih.gov/geo), ArrayExpress (http://www.ebi.ac.uk/arrayexpress), and the UCSC Cancer Genomics Browser (https://genome-cancer.ucsc.edu). This collection contains datasets of expression profiles of benign prostate tissue, primary tumors, and metastatic or CRPC.

To assess the association of CCM1 gene expression with OS or metastasis-free survival, clinical information was extracted from the Swedish Watchful Waiting Cohort and Johns Hopkins Cohort. Patients in each cohort were subdivided into categories of “low” (<50th percentile) or “high” (≥50th percentile) CCM1 expression. Kaplan–Meier curves for OS and metastasis-free survival were drawn for each category, and Cox proportional hazard regression analysis was performed for the statistical comparison of survival rates between low and high CCM1 expression groups.

We examined the presence of copy number alterations and genomic rearrangements in the vicinity of the CCM1 and DDX5 genes based on the processed variant calls from the previous studies including the Canadian prostate cancer genome network (31-33). We quantified the amplification events of these genes by counting the number of samples that have copy number gain of the genomic segments containing CCM1 or DDX5 genes. We also investigated genomic rearrangements that might affect the expression level of these genes in the genomic region including 1 Mb on both sides of the respective genes.

## Table Legends

**Table S1.**
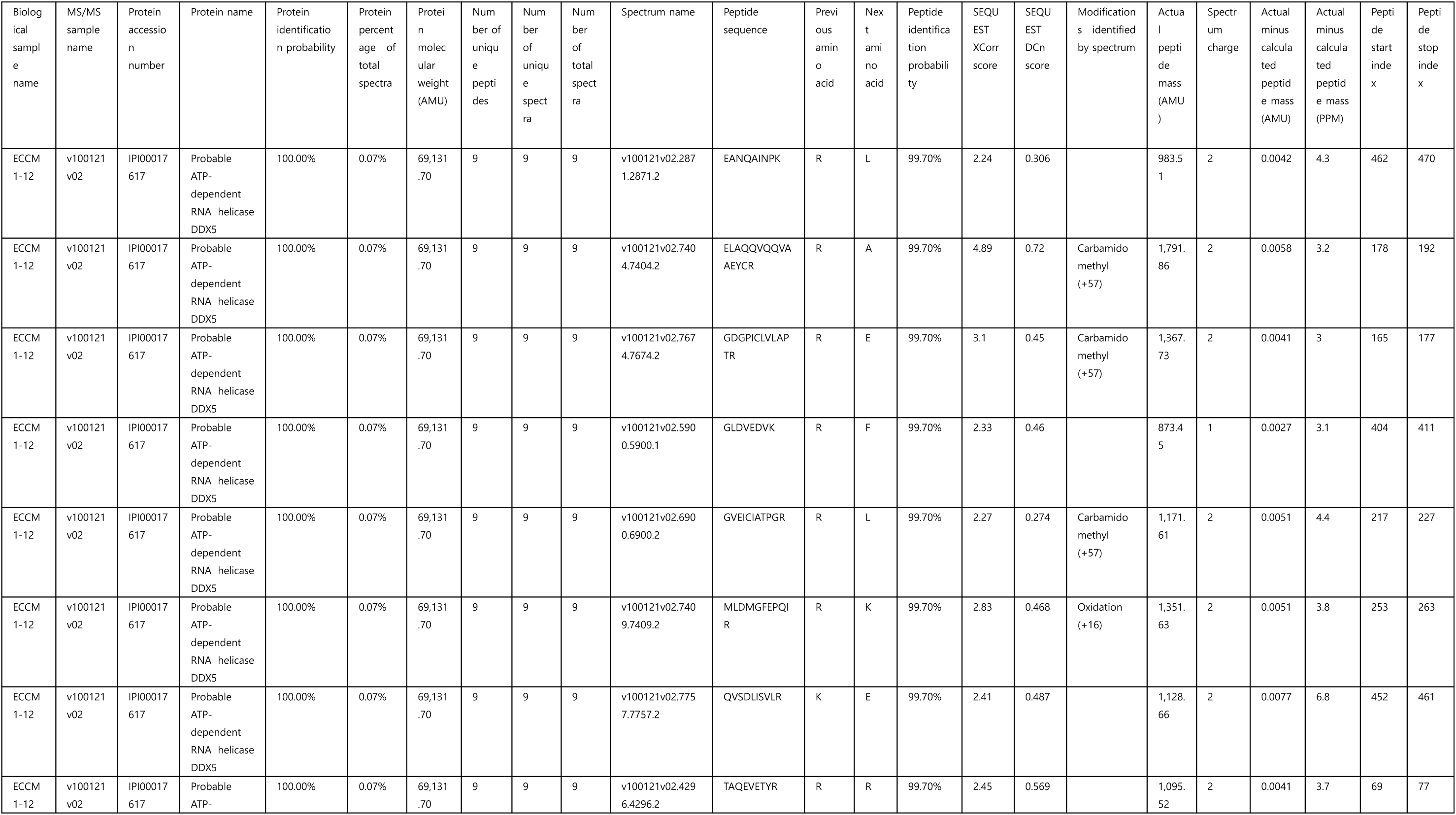
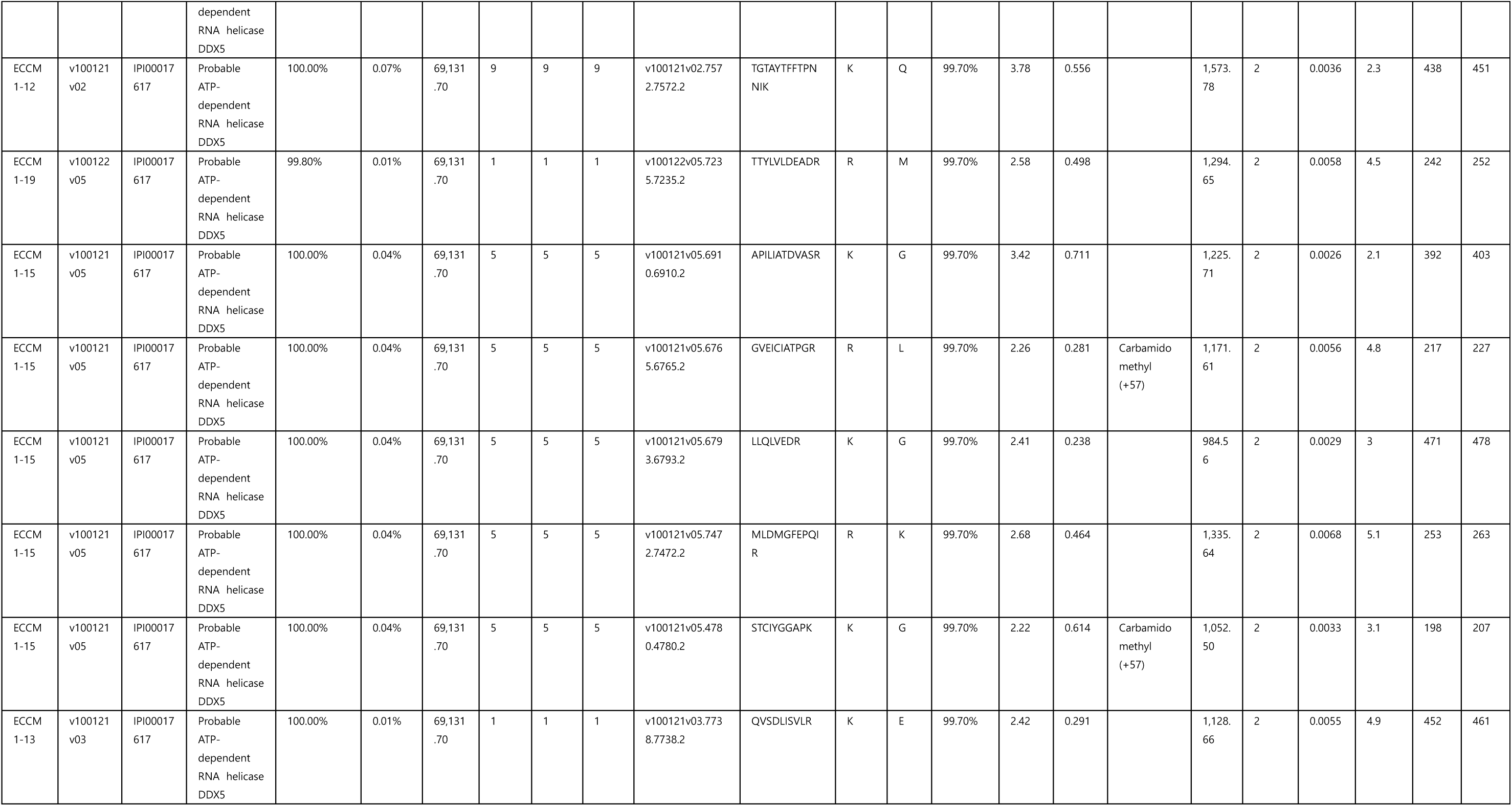
Proteomics identification of DDX5 from a pull-down experiment. The biological sample names indicate specific position of gel slices from an SDS-PAGE gel.

## Figure Legends

**Figure S1.**
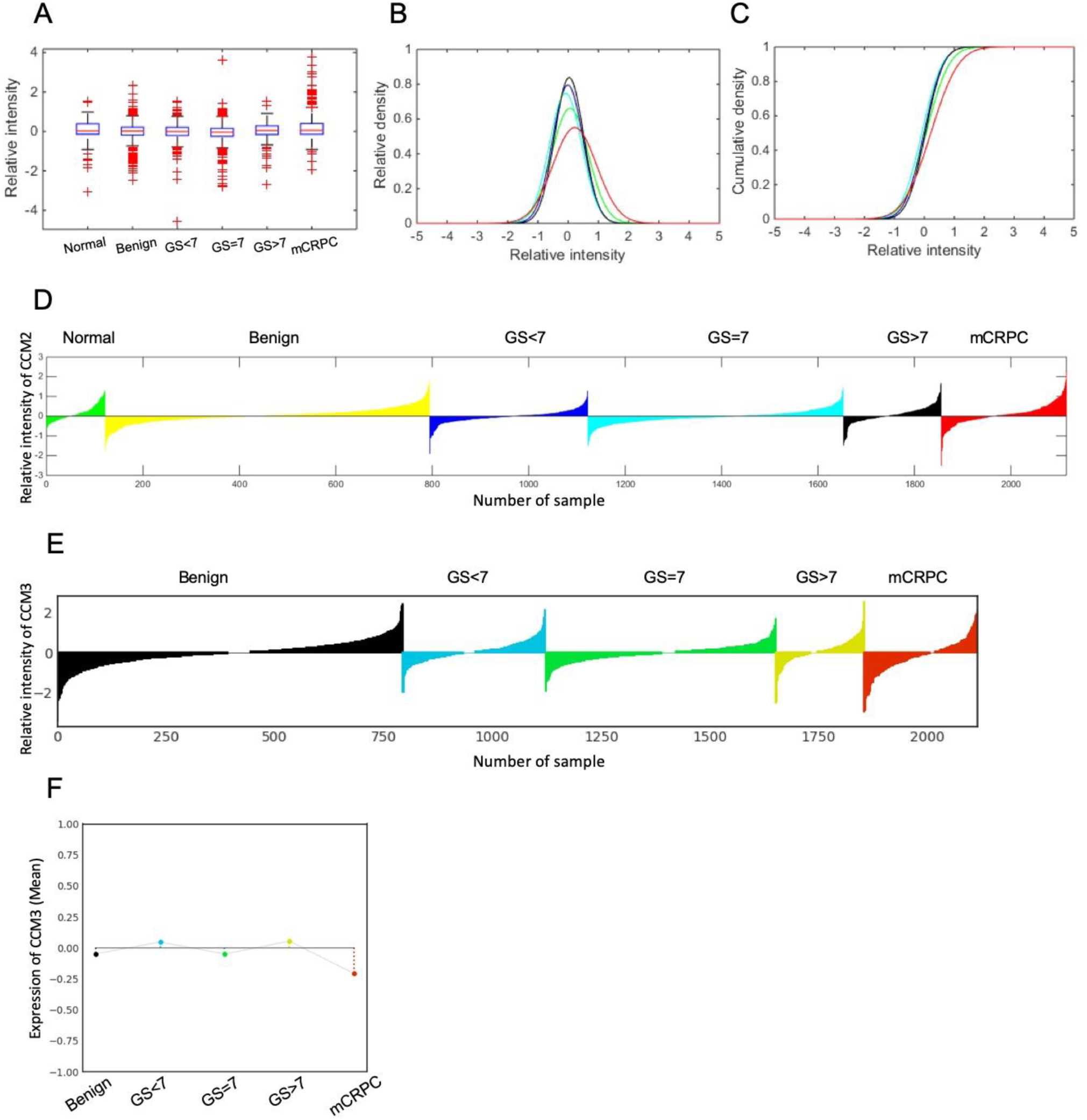
Expression analysis of CCM1, CCM2, and CCM3. (A) Boxplot of CCM1 gene expression in different disease states. The x-axis presents the disease state, and the y-axis presents normalized gene expression (log2 scale). (B) The distribution of samples in each disease state according to their relative CCM1 gene expression. The x-axis presents normalized gene expression, and the y-axis presents the relative density of the samples. (C) The cumulative density of the samples according to their CCM1 gene expression levels. The x-axis presents normalized gene expression, and the y-axis presents the cumulative density of the samples. Different line colors in panels B–C indicate different diseases as described for Figure 1C. (D) The waterfall plot displays the normalized CCM2 gene expression of individual samples from the prostate cancer transcriptome atlas (PCTA) cohort. (E) This waterfall plot displays the normalized CCM3 gene expression of individual samples from the PCTA cohort. (F) The stem plots with different colors show the mean expression of CCM3 in each disease state. The y-axis presents normalized gene expression (log2 scale).

**Figure S2.**
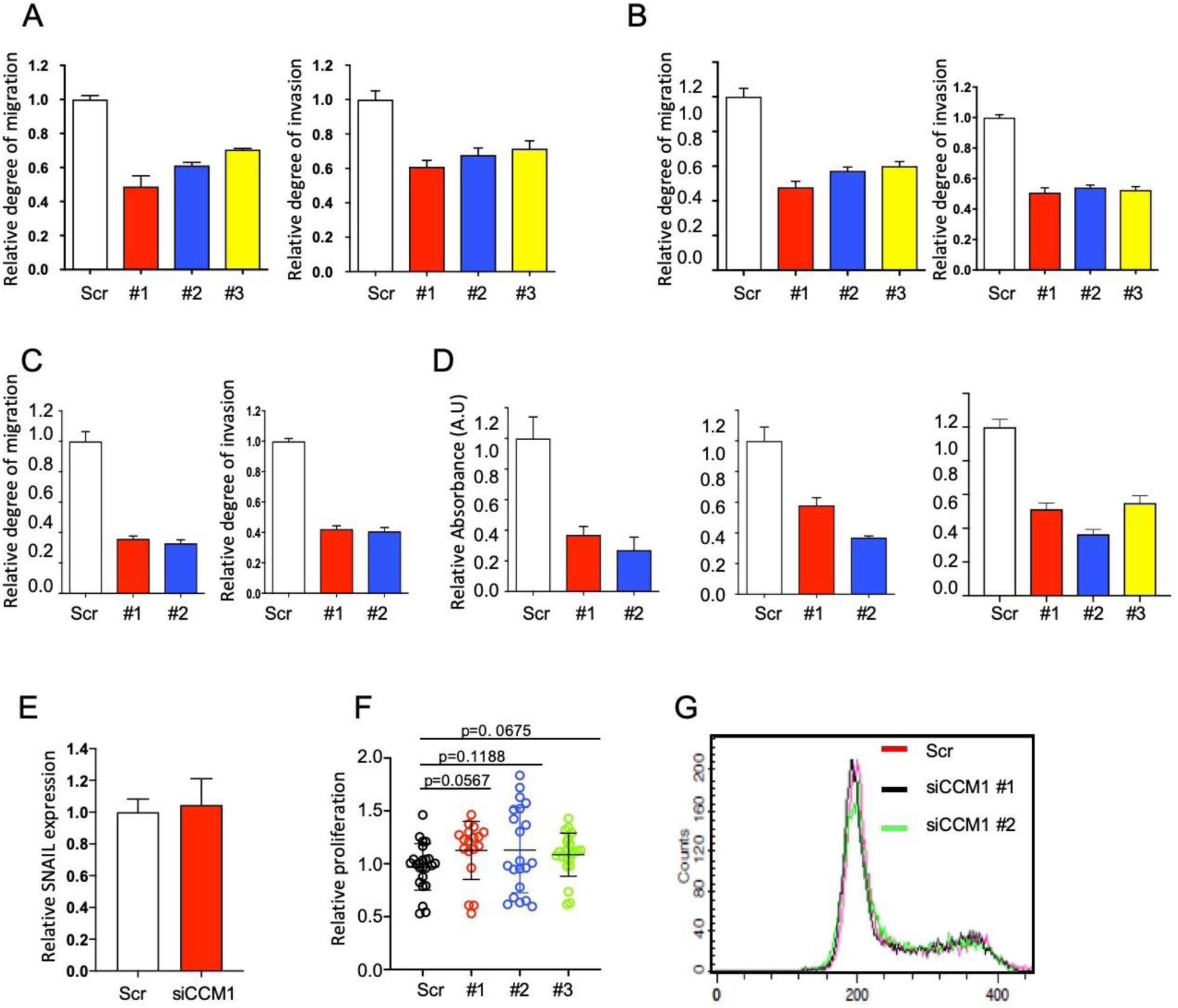
Suppression of CCM1 represses a metastatic hallmark in prostate cancer cells *in vitro*. (A) DU145, (B) C4-2, and (C) CWR22r shCCM1 cells were plated on Transwell membranes coated without or with Matrigel for migration (left) and invasion (right) assays, respectively, incubated for up to 72 h, and stained. The mean ± SEM is presented as a ratio to the expression in the scramble (Scr) control. (D) LNCaP (left), C4-2 (middle), and CWR22r (right) cells were grown in soft agar for 7 days, and each lysate was analyzed using a microplate reader to measure anchorage-independent survival. The data are presented as the mean ± SEM of absorbance expressed as a ratio to the expression in the Scr control. (E) SNAIL expression was analyzed via qPCR in CCM1-suppressed PC3 cells. (F) LNCaP shCCM1 cells were plated, grown for 3 days, and cellular proliferation was analyzed using Ez-Cytox assay kits. Scatter plots present the ratio of endpoint values (n = 3– 5/set) from five independent experimental sets relative to control means in each corresponding experiment. Welch’s unpaired *t*-test was used for statistical analysis. (G) CCM1 was silenced in DU145 cells using two different siRNAs, and cell cycle progression was analyzed.

**Figure S3.**
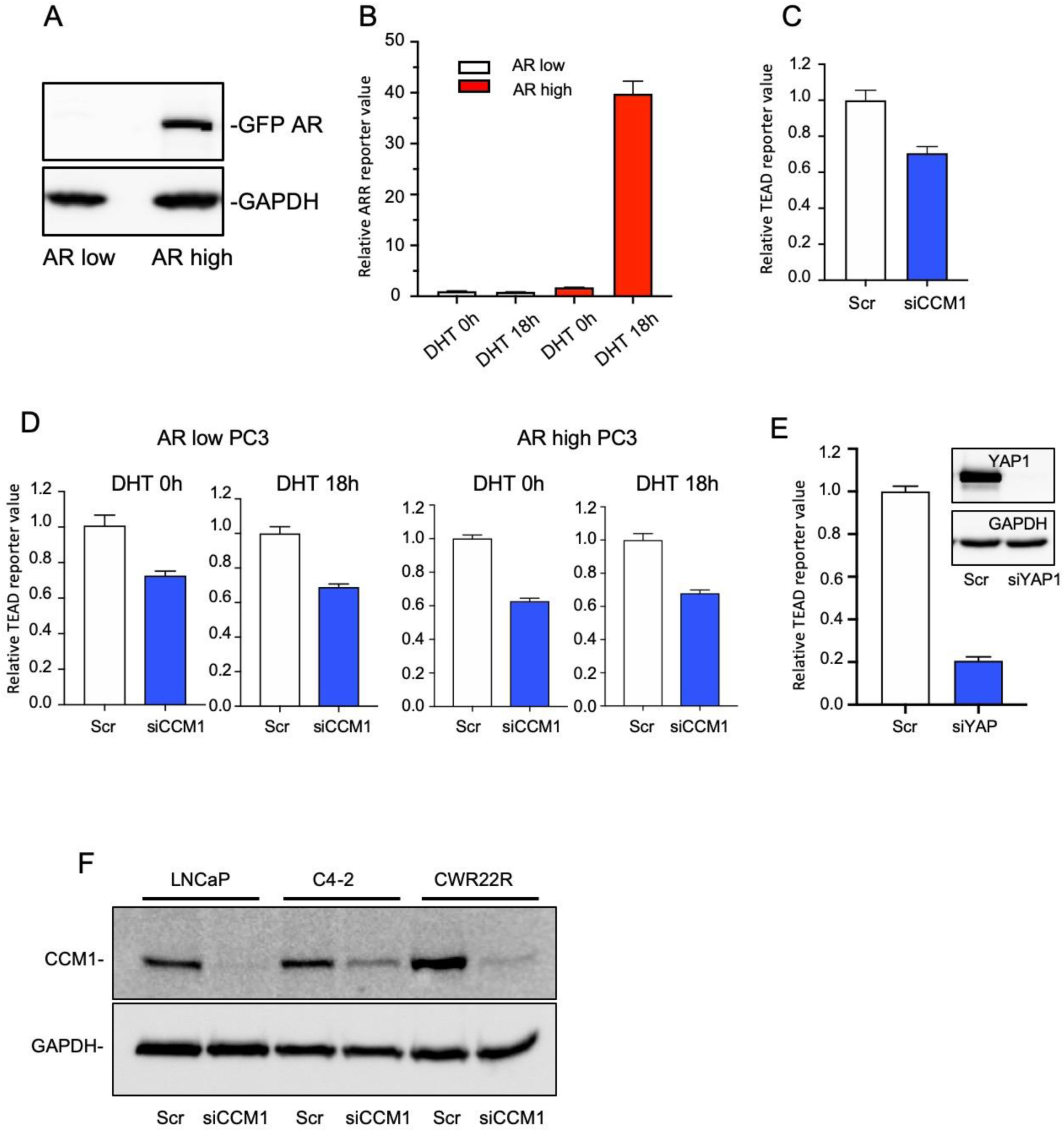
CCM1-mediated regulation of Yes-associated protein (YAP)/transcriptional co-activator with PDZ-binding motif (TAZ) signaling. (A) An androgen receptor (AR) overexpression plasmid was stably transfected in WT PC3 cells, and individual PC3 clones were sorted into AR high and AR low groups according to the expression level of ectopic AR. (B) PC3 AR high (red box) or low (white box) cells were grown in medium supplemented with 10% charcoal-stripped FBS and stimulated with 1 µM DHT for the indicated time points, and ARR reporter activities were measured using luciferase assays. (C) CCM1 was RNAi-silenced in DU145 cells grown in medium containing 10% FBS, and TEAD reporter activity was analyzed. (D) CCM1 was RNAi-silenced in AR low (left two panels) or AR high (right two panels) PC3 cells grown in 10% charcoal-stripped FBS supplemented medium with or without DHT stimulation as indicated, and TEAD reporter activity was analyzed. Reporter values are presented relative to that in each scramble control. (E) YAP1 was suppressed via RNAi in LNCaP cells, and TEAD reporter activity was analyzed. Suppression of YAP1 was confirmed via immunoblotting (Inset). (F) The representative degree of CCM1 suppression in LNCaP, C4-2, and CWR22r cells.

**Figure S4.**
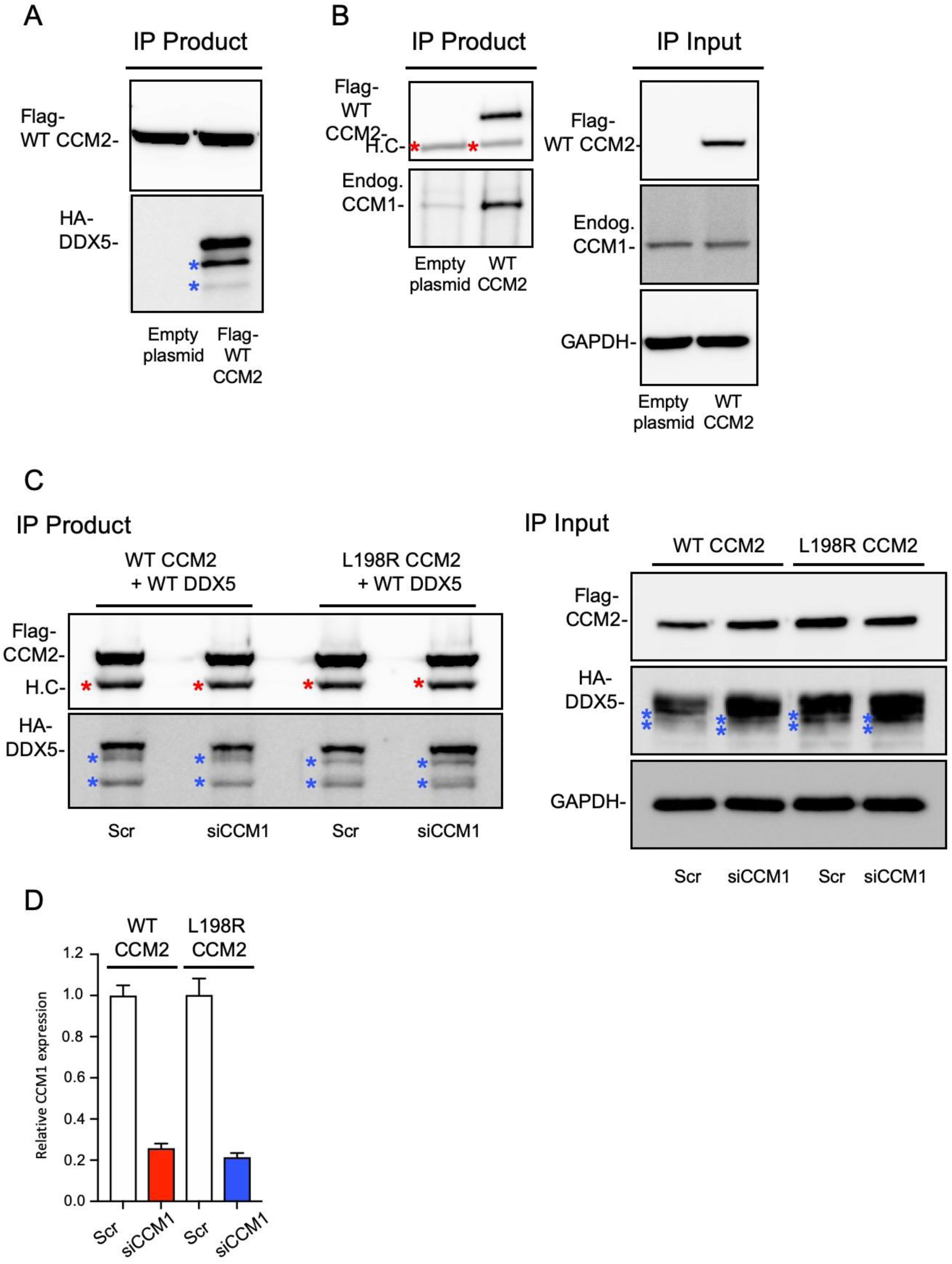
DDX5 interacts with CCM complexes. (A) HA-tagged DDX5 and either Flag-tagged WT CCM2 or L198R CCM2 were co-expressed in PC3 cells. Flag-CCM2 were immunoprecipitated with Flag antibody, and co-immunoprecipitated DDX5 was immunoblotted with anti-Flag or anti-HA antibody. (B) Flag-tagged WT CCM2 was overexpressed in PC3 cells and immunoprecipitated with anti-Flag antibody. Co-immunoprecipitated endogenous CCM1 was immunoblotted using anti-CCM1 antibody. (C) Shown is another replicated experiment performed as described in Figure 4C. HA-DDX5 and Flag-tagged WT or L198R CCM2 was co-overexpressed in PC3 cells and immunoprecipitated with anti-Flag antibody, and co-immunoprecipitated DDX5 was immunoblotted with anti-HA antibody (left panel). Expression levels of overexpressed CCM2 and DDX5 in immunoprecipitation input samples are shown (right panel). RNAi silencing of CCM1 was performed in CCM2-overexpressing samples as indicated. Red asterisks: heavy chain, blue asterisks: degradation products. (D) Suppression of CCM1 in panel C was additionally analyzed via qPCR.

**Figure S5.**
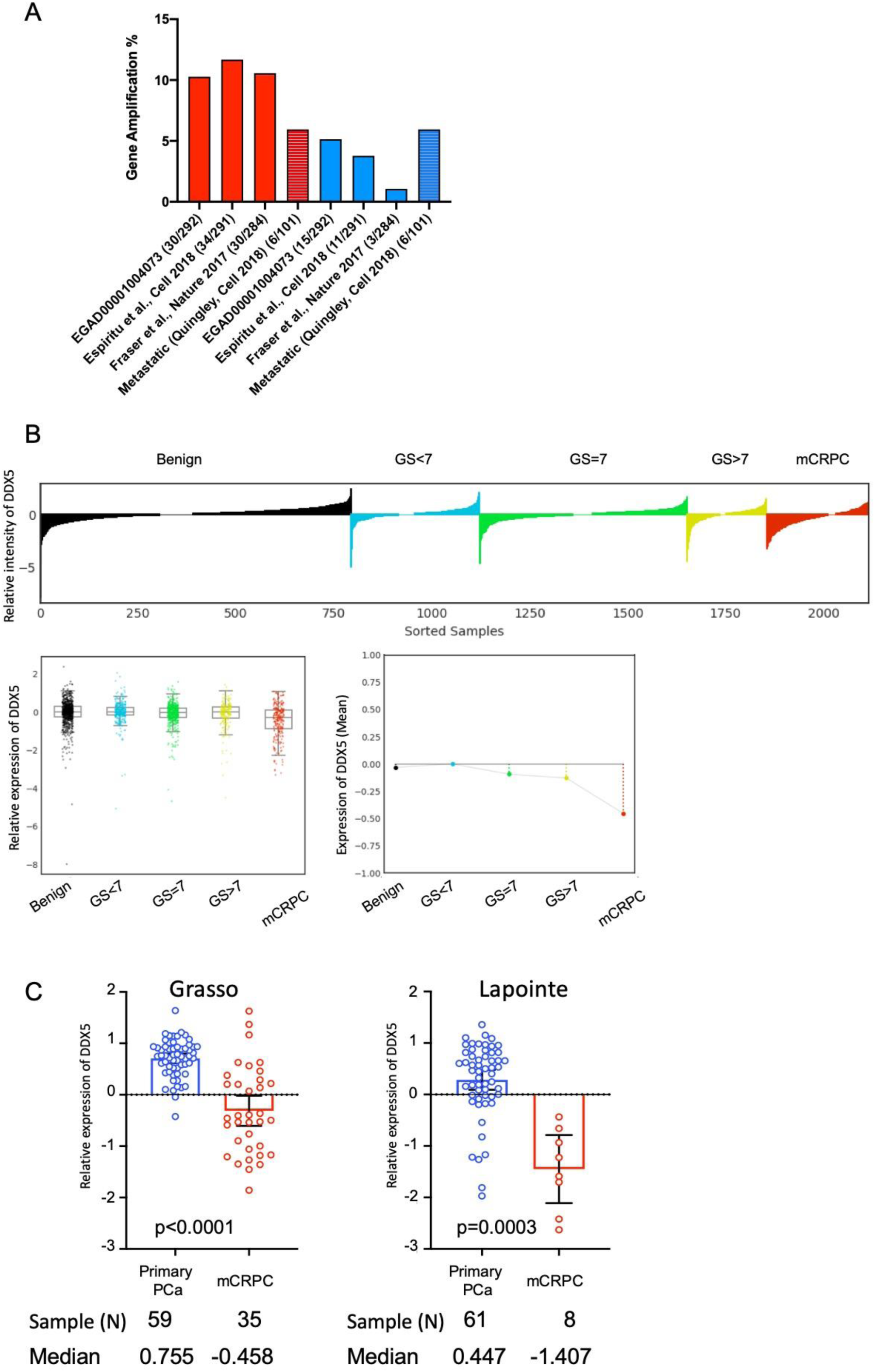
Regulatory mechanisms of CCM1 and DDX5 expression. (A) Shown are comparisons of the frequencies of the gene amplification of CCM1 (red bars) and DDX5 (blue bars) from four independent cohort studies. In each group of four bars, the first three bars represent data from primary prostate cancer studies, and the fourth bar represents data from a metastatic castration-resistant prostate cancer (mCRPC) study. The y-axis presents percentiles (100 × [number of samples with gene amplification/number of total samples]). (B) The waterfall plot displays normalized DDX5 gene expression (log2 scale) for individual samples from the prostate cancer transcriptome atlas cohort (top). The Wilcoxon rank sum test results between subsets (mCRPC vs. primary PCa): fold change = −0.383, P < 0.001. (bottom left) Boxplot of DDX5 gene expression in different disease states. The x-axis presents the disease state, and the y-axis presents normalized gene expression. (bottom right) The stem plots with different colors show the mean expression levels of DDX5 in each disease state. The y-axis values represent normalized expression of DDX5. (C) Scatter plot analysis of the relative expression of DDX5 between primary PCa and metastatic CRPC samples from two transcriptome studies (Grasso (35) and Lapointe (36) studies). Y-axis values were, which were retrieved from the Oncomine database, are shown as the log2 median-centered ratio. Mean with 95% confidence intervals. Welch’s unpaired *t*-test was used for statistical significance.

**Figure S6.**
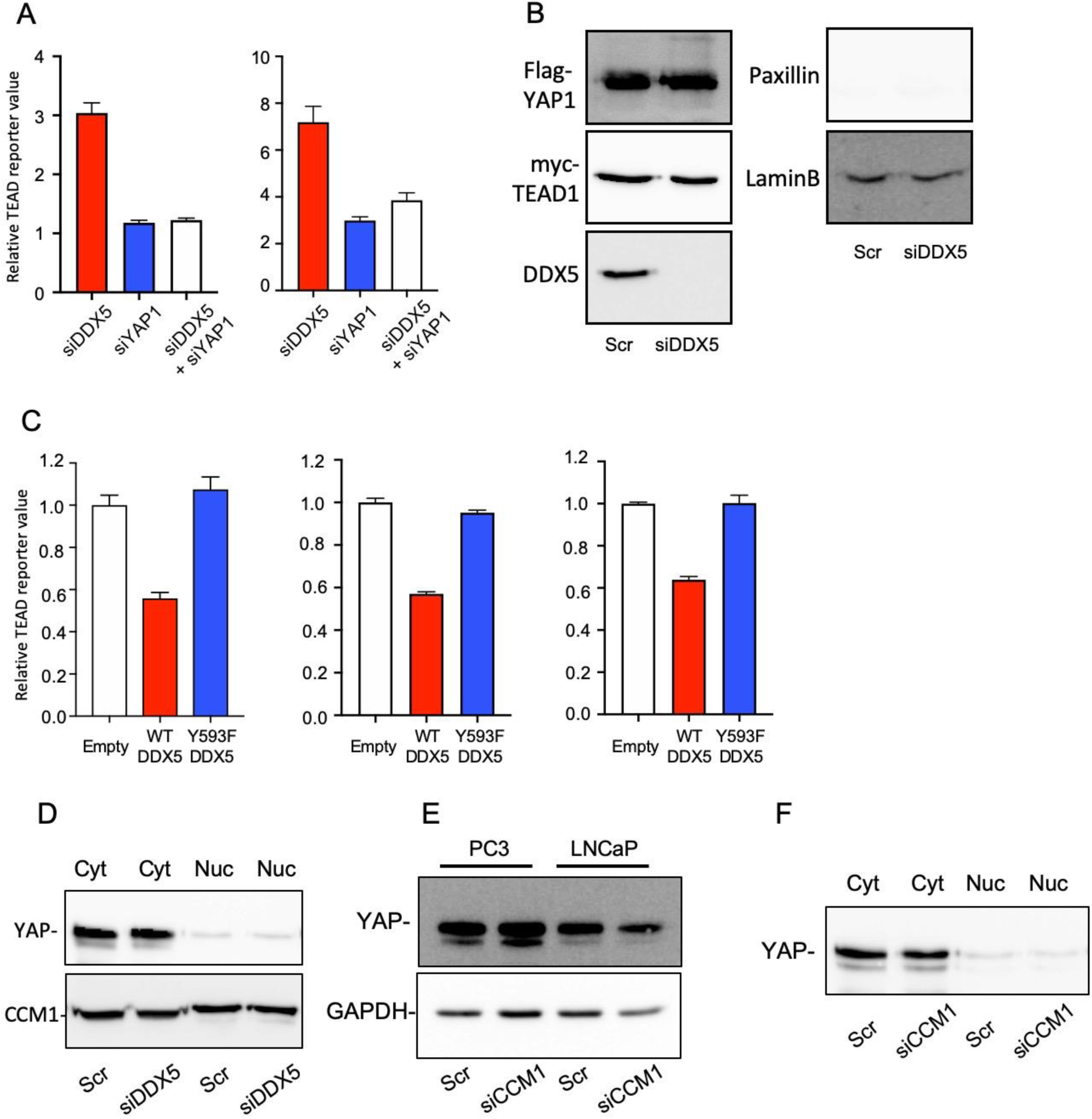
CCM1 and DDX5 regulate Yes-associated protein (YAP)/transcriptional co-activator with PDZ-binding motif signaling. (A) RNAi-mediated suppression of DDX5 or YAP1 alone or co-suppression of DDX5 and YAP1 was performed, and TEAD reporter activity was analyzed in C4-2 (left) and C4-2B (right) cells grown in medium supplemented with charcoal-stripped FBS and stimulated with DHT for 18 h. (B) PC3 cells co-transfected with overexpression plasmids for Flag-YAP, myc-TEAD1, and siDDX5, were harvested, and nuclear lysates were generated. Shown are immunoprecipitation input samples from the nuclear lysates used in Figure 5H. Paxillin, a cytosolic marker, is not shown in these nuclear lysates. (C) WT or Y593F DDX5 was overexpressed in LNCaP (left), C4-2B (middle), and CWR22r (right) cells, and TEAD reporter activity was measured. (D) Subcellular localization of YAP was analyzed in DDX5-suppressed PC3 cells. (E) YAP expression was analyzed in the total lysates of CCM1-suppressed PC3 and LNCaP cells. (F) Subcellular localization of YAP was analyzed in CCM1-suppressed PC3 cells. Cyt: cytosolic fraction, Nuc: nuclear fraction.

**Figure S7.**
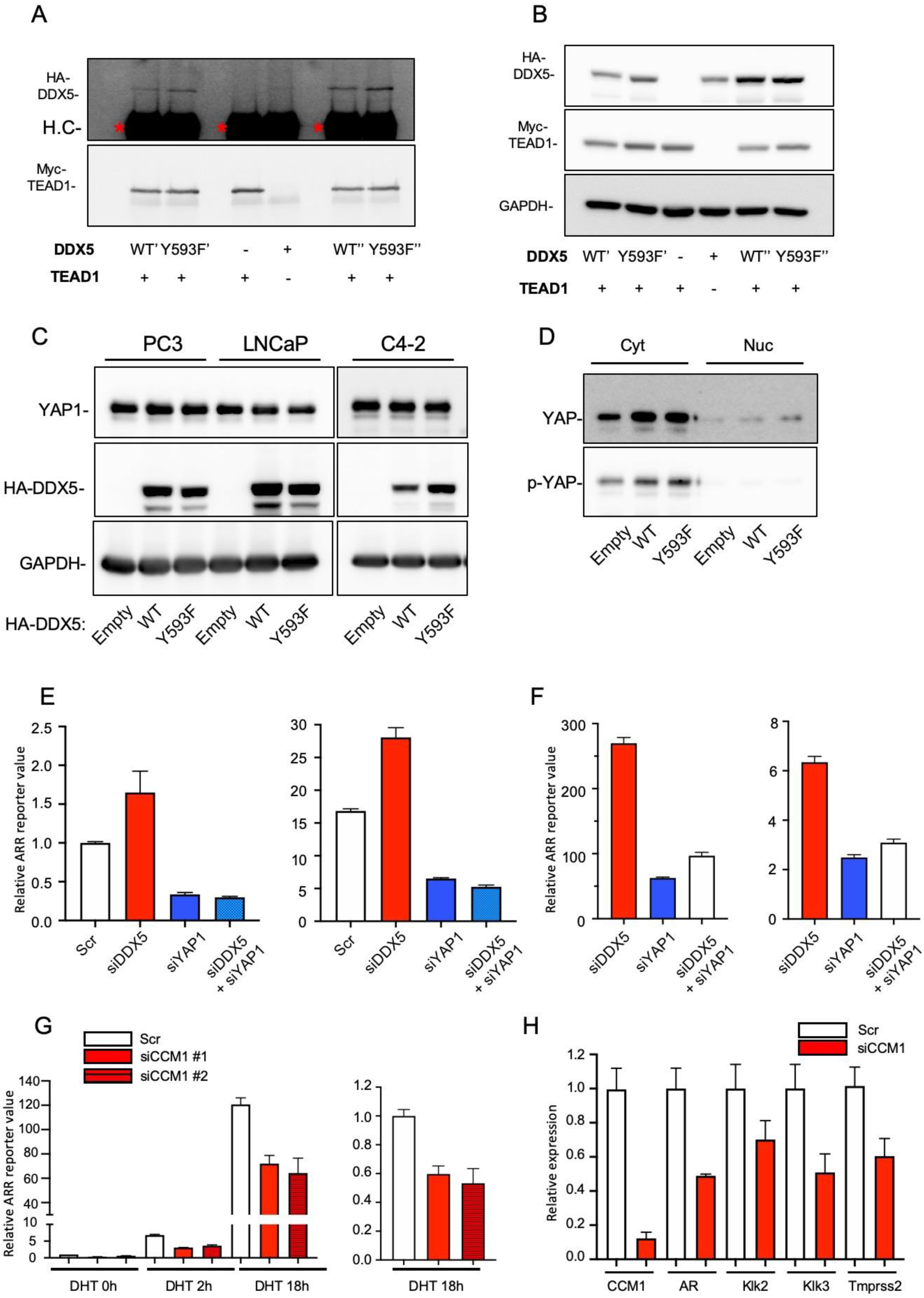
Regulatory mechanisms of DDX5 and CCM1 in Yes-associated protein (YAP)/transcriptional co-activator with PDZ-binding motif (TAZ) and androgen receptor (AR) signaling. (A) HA-tagged WT or Y593F DDX5 and myc-tagged TEAD1 were co-expressed in C4-2 cells, the total lysates were immunoprecipitated with anti-myc antibodies, and immunoblotted with anti-myc or anti-HA antibodies to detect immunoprecipitated myc-TEAD1 and co-immunoprecipitated HA-DDX5, respectively. Our co-immunoprecipitation assays with DDX5-overexpressing lysates were replicated as shown (′, ″). (B) Shown are the immunoprecipitation input samples of panel (A). (C) DDX5 was overexpressed in PC3, LNCaP, and C4-2 cells, and YAP1 levels in each total lysate were analyzed via immunoblotting. (D) Cytosolic and nuclear fractions of C4-2 cells expressing ectopic WT or Y593F DDX5 were separated, and the endogenous levels of total YAP and S127-phosphorylated YAP in cytosolic and nuclear fractions were analyzed. Cyt: cytosolic fraction, Nuc: nuclear fraction. (E, F) RNAi-mediated silencing of DDX5 or YAP1 or co-silencing of DDX5 and YAP1 and transfection of ARR reporter plasmids were performed using PCa cells grown in charcoal-stripped FBS-supplemented medium and stimulated with DHT for 18 h. (E) LNCaP cells were stimulated without (left) or with 1 µM DHT (right), and ARR reporter activity was analyzed. (F) C4-2 (left) or C4-2B (right) cells were stimulated with DHT, and ARR reporter activity was analyzed. The data are presented as a mean ± SEM relative to each untreated scramble (Scr) control. Graphs were generated from three independent experiments. (G) LNCaP cells were co-transfected with two different CCM1 siRNAs (#1, #2) and ARR reporter plasmids, grown in charcoal-stripped FBS-supplemented medium, and stimulated with 1 µM DHT for the indicated times. Reporter values are presented relative to those in the Scr control without DHT stimulation (left). Ratio of reporter activities after 18 h of DHT treatment relative to those in the Scr control (right). (H) CCM1 was suppressed in LNCaP cells, and the expression of CCM1, androgen receptor (AR), and representative AR target genes was analyzed via qPCR.

